# Nuanced role for dendritic cell intrinsic IRE1 RNase in the regulation of antitumor adaptive immunity

**DOI:** 10.1101/2022.07.20.500838

**Authors:** Felipe Flores-Santibañez, Sofie Rennen, Dominique Fernandez, Clint De Nolf, Sandra Gaete, Camila Fuentes, Carolina Moreno, Diego Figueroa, Álvaro Lladser, Takao Iwawaki, María Rosa Bono, Sophie Janssens, Fabiola Osorio

**Author notes:** These authors equally contributed to this work (Felipe Flores-Santibañez, Sofie Rennen). Corresponding authors. **Correspondence:** Fabiola Osorio, Immunology Program, Institute of Biomedical Sciences, Faculty of Medicine, Universidad de Chile, Av. Independencia 1027, Postal code: 8380453, Santiago, Chile, Sophie Janssens, Laboratory for Endoplasmic Reticulum Stress and Inflammation VIB Center for Inflammation Research, Technologiepark 71, B-9052 Zwijnaarde, Ghent, Belgium.

## Abstract

The IRE1/XBP1s axis of the unfolded protein response (UPR) plays divergent roles in dendritic cell (DC) biology in steady state versus tumor contexts. Whereas tumor associated DCs show dysfunctional IRE1/XBP1s activation that curtails their function, the homeostasis of conventional type 1 DCs (cDC1) in tissues requires intact IRE1 RNase activity. Considering that cDC1s are key orchestrators of antitumor immunity, it is relevant to understand the functional versus dysfunctional roles of IRE1/XBP1s in tumor DC subtypes. Here, we show that cDC1s constitutively activate IRE1 RNase within subcutaneous B16 melanoma and MC38 adenocarcinoma tumor models. Mice lacking XBP1s in DCs display increased melanoma tumor growth, reduced T cell effector responses and accumulation of terminal exhausted CD8^+^ T cells. Transcriptomic studies revealed that XBP1 deficiency in tumor cDC1s decreased expression of mRNAs encoding XBPls and regulated IRE1 dependent decay (RIDD) targets. Finally, we find that the dysregulated melanoma growth and impaired T cell immunity noticed in XBP1 deficient mice are attributed to RIDD induction in DCs. This work indicates that IRE1 RNase activity in melanoma/MC38-associated DCs fine tunes aspects of antitumor immunity independently of XBP1s, revealing a differential role for the UPR axis that depends on the DC subtype and cancer model.

## INTRODUCTION

A crucial arm of antitumor immunity relies on effective activation of tumor specific cytotoxic CD8^+^ T cells endowed with the ability to eliminate cancer cells. This process is critically dependent on type 1 conventional dendritic cells (cDC1), which excel in cross-presentation of tumor-associated antigens (1–3), secrete soluble factors that potentiate CD8^+^ T cell function (2, 4–7), and prevent the generation of terminal exhausted CD8^+^ T cells committed to irreversible dysfunctional phenotypes in tumors (8). Tumor cDC1 infiltration correlates with improved prognosis (2, 5, 9) and better response to immune checkpoint blockade (ICB) therapy (9–12). Besides, therapeutic strategies expanding/activating tumor cDC1s have shown promising results in clinical trials (11, 13, 14). Additional DC subsets such as type 2 DCs (cDC2s), and a novel DC activation state termed ‘mregDCs’ (mature DCs enriched in immunoregulatory molecules) can also boost antitumor CD4^+^ and CD8^+^ T cell responses (7, 15–17), indicating that cDCs are interesting candidates in immunotherapy. However, the molecular mechanisms safeguarding the function of these cells in tumors have not been fully elucidated.

An emerging intracellular pathway regulating DC biology is the inositol-requiring enzyme 1 alpha (IRE1) branch of the unfolded protein response (UPR), which is an adaptive cellular response maintaining the fidelity of the cellular proteome (18). Upon endoplasmic reticulum (ER) stress, the endoribonuclease (RNase) domain of IRE1 splices *Xbp1* mRNA, generating the transcription factor XBP1 spliced (XBP1s), master regulator of protein folding and ER biogenesis (18–20). The IRE1 RNase domain can also promote the degradation of a subset of mRNAs/miRNAs in a process known as ‘regulated IRE1-dependent decay’ (RIDD) (21), which is a mechanism beginning to be understood in pathological settings including metabolism, inflammation and cancer (22–24).

In steady state, IRE1 regulates cDC homeostasis via constitutive activation of its RNase domain, a feature noticed in cDC1s across several tissues (25, 26). Furthermore, cDC1s are markedly sensitive to perturbations in the IRE1/XBP1s axis, as genetic loss of the transcription factor XBP1 alters proteostatic programs and counter activates the RIDD branch, which mediates the decay of various mRNAs involved in integrin expression, ER to golgi transport and antigen presentation, among others (25, 26). The selectivity of the IRE1/XBP1s axis in cDC1s is underscored in microarray studies of XBP1 deficient cells, which change the transcriptomic landscape of cDC1s but not cDC2s (26). As such, cDC1s opt the IRE1/XBP1s axis for proper function in steady state, but it is unclear if the pathway displays similar roles in cDC1s infiltrating tumors. This is a relevant question, as reported work shows that metabolically stressed tumors elicit maladaptive UPR activation in certain tumor immune cells (including DCs), which reprograms their phenotype towards dysfunctional states that promote tumor growth (27, 28). For instance, DCs infiltrating ovarian cancer (typified by expression of the cDC2/monocyte marker CD11b^+^ (29)) display persistent IRE1/XBP1s activation that triggers aberrant intracellular lipid accumulation, resulting in impaired immunostimulatory functions and leading to tumor progression (30). Thus, these data suggest that IRE1 may play different, or even opposite roles in DC biology depending on the subtype or the inflammatory context. As such, a correct delineation of the role of the enzyme in tumor cDCs is required to understand if intervention of this UPR branch can be targeted for potential cancer therapies.

In this work, we study the role of IRE1/XBP1s in DCs by focusing on two immunoresponsive tumor models: subcutaneous mouse B16/B78 melanoma and MC38 colon adenocarcinoma (2, 10). We identified that cDC1s display constitutive IRE1 RNase activity in tumors, which follows a lineage-intrinsic trait not influenced by the tumor microenvironment. In contrast to previous reports (30), deletion of XBP1 in DCs did not decrease tumor burden. Rather, XBP1 deficiency in DCs resulted in a moderate increase of tumor growth, lower frequencies of intratumoral effector T cells, and accumulation of terminal exhausted TIM-3^+^CD8^+^ cells in the melanoma model. Transcriptomic studies revealed that XBP1 deficient tumor cDC1s downregulate proteostatic processes and decrease the expression of mRNAs encoding XBP1s and regulated IRE1 dependent decay (RIDD) targets. Importantly, animals bearing double deletion of IRE1 RNase and XBP1 in DCs display normal tumor growth and adaptive immunity in the melanoma model, highlighting a role for IRE1 RNase hyperactivation in fine tuning aspects of antitumor immunity via DCs.

## RESULTS

### cDC1s constitutively activate IRE1 RNase in subcutaneous melanoma and MC38 colon carcinoma tumors

The tumor microenvironment contain activators of the IRE1/XBP1s axis that are detected by immune cells (27, 30–32). To identify relevant cell types activating IRE1 RNase in tumors, we analyzed the immune composition of B16 tumors of ERAI mice, a mouse strain that reports IRE1 RNase activity through expression of Venus Fluorescent Protein (VenusFP) fused with the sequence of *Xbp1s* mRNA (33) (validated in (25, 26, 31)). To allow unsupervised identification of IRE1 RNase cellular targets, we devised a 17-color flow cytometry panel and data was visualized on a *t*-distributed stochastic neighbor embedding (*t*-SNE) map. Cells were grouped into populations by DBScan-guided automated clustering (Fig. 1A-B, Supp. Fig. 1A), which identified 15 cell clusters that included CD8^+^ T cells, CD4^+^ T cells, monocyte-derived cells (MdCs), MHC-II-expressing MdCs, NK cells, NKT cells, B cells, neutrophils and cDC1s. As expected (17), cDC2s and tumor associated macrophages (TAMs) (clusters 4 and 6) showed a degree of heterogeneity and convergence. Also, our analysis revealed two undefined clusters based on surface marker expression; Cluster 11: CD4^+^ CD11c^+^ CD26^+^, and Cluster 14: CD3^+^ CD4^+^CD11b^int^ F4/80^+^ MHC-II^high^ CD11c^int^ CD26^high^.

**Figure 1.**
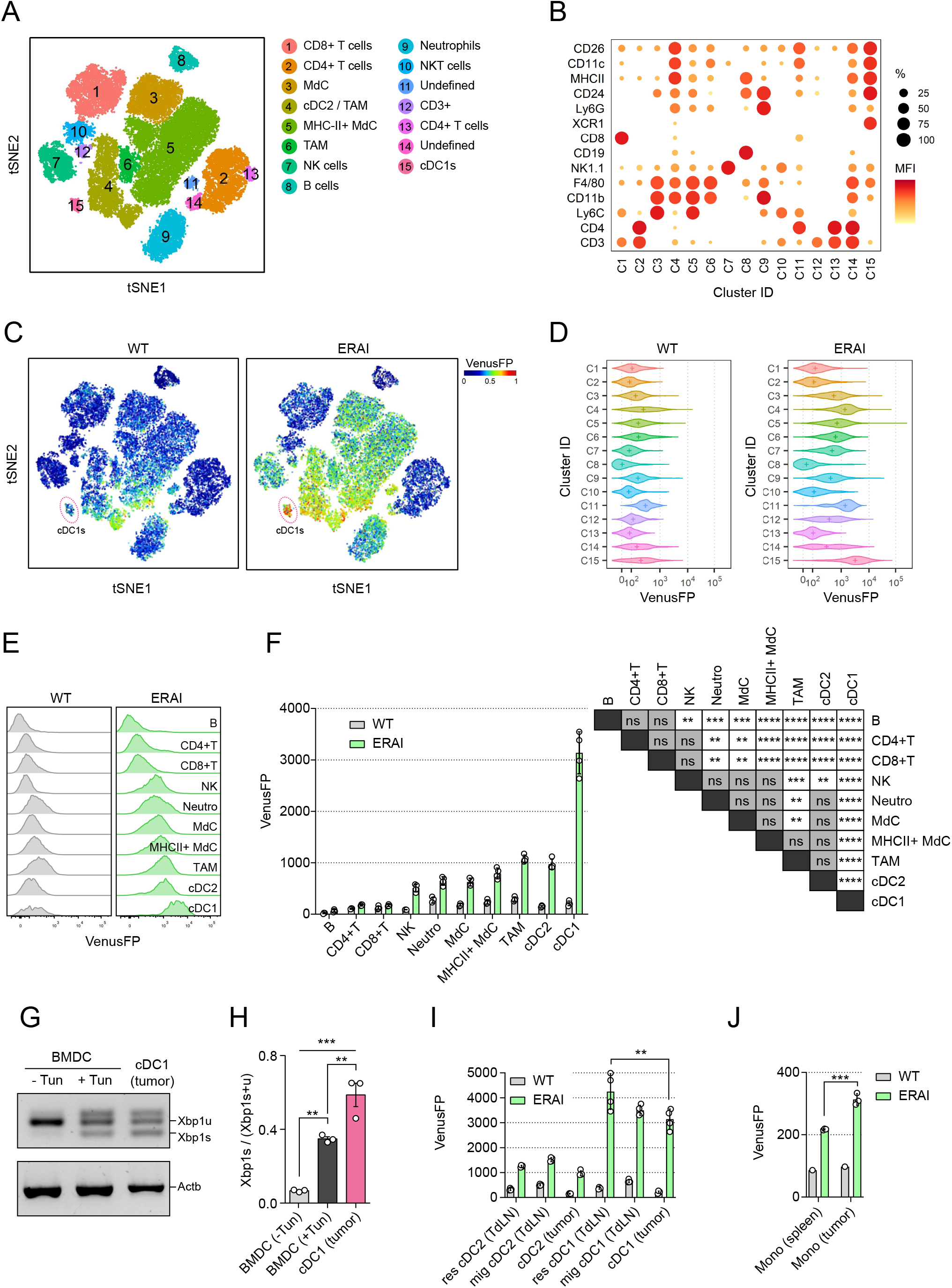
cDC1s are predominant cellular targets of IRE1 activation in melanoma tumors. **(A-F)** B16-F10 melanoma cells were implanted intradermally on ERAI or control mice and 11 days after implantation, tumor tissue was analyzed by multicolor flow cytometry. n=4 mice per group, representative of two independent experiments. **(A)** t-SNE of 40.000 immune (CD45+) infiltrating cells from melanoma of ERAI or control mice. Colors indicate unsupervised clustering by DBSCAN. **(B)** Marker expression across the different cell clusters identified in (A). See also Supp. Fig 1A. **(C)** t-SNE map colored by VenusFP signal intensity from control or ERAI mice. cDC1 cluster is highlighted in a red circle. **(D)** VenusFP signal quantification across the different cell clusters identified in (A). Median fluorescence intensity for VenusFP is depicted with a “+” inside each violin plot. **(E-F)** Histograms (E) and quantification (F) of VenusFP signal from manually gated immune populations from B16-F10-bearing WT and ERAI mice (see gating strategy in Supp. Fig.2A). Statistical significance is depicted as compact letter display, ANOVA and Tukey post-test between ERAI mice. **(G-H)** Xbp1 splicing assay (G) of tumor cDC1s isolated by cell sorting from B16-FLT3L-bearing WT mice compared to BMDCs treated with 1 μg/mL of the ER-stressor Tunicamycin (Tun) or medium for 8h. (H) Bars show the image quantification of the ratio between Xbp1 spliced (Xbp1s) and total Xbp1 (Xbp1s + Xbp1u). n= 3 samples per group (representative of three independent experiments), mean ± s.e.m, ANOVA and Tukey post-test, ** p<0.01, *** p<0.001. **(I)** Quantification of VenusFP signal from tumor and tumor draining lymph node migratory (mig) and resident (res) cDCs (see gating strategy in Supp. Fig.2B). ** p<0.01, ANOVA and Tukey post-test between ERAI mice. n=4 mice per group, representative of two independent experiments, mean ± s.e.m. **(J)** Quantification of VenusFP signal from intratumoral and spleen monocytes (CD11b^hi^ Ly6C^hi^ cells). *** p<0.001, t-test between ERAI mice. n= 3 mice (ERAI) or 1 mouse (WT), representative of two independent experiments, mean ± s.e.m.

Next, IRE1 RNase activity from the ERAI reporter mice line was determined in the clusters of the t-SNE plot (Fig. 1C-D). Data indicated that among CD45^+^ hematopoietic cells, cDC1s represented the population with highest fluorescence intensity of VenusFP (Fig 1C-D, cluster 15). Additional immune cells including cDC2s, MdC, MHC-II^+^MdC, TAM, neutrophils, NK cells and cells from cluster 11 also showed noticeable VenusFP induction, albeit at lower levels than cDC1s; whereas CD4^+^ T cells, CD8^+^ T cells and B cells showed little or no induction of VenusFP compared to cells from control animals (Fig 1D). Manual gating analysis (Fig. 1E-F, Supp. Fig. 2A) confirmed that the mean fluorescence intensity (MFI) of VenusFP in melanoma-associated cDC1s was higher than additional myeloid and lymphoid cells (Fig. 1E-F). Notably, similar results were observed in cDC1s infiltrating MC38 tumors (Supp. Fig. 1B). Finally, these data was confirmed by PCR for endogenous *Xbp1* spliced and unspliced forms from tumor cDC1s isolated from wild type (WT) animals implanted with the Bl6-FLT3L melanoma cell line (which expresses the DC-differentiation factor FMS-like tyrosine kinase 3 ligand (FLT3L) (11, 34)). Data in Fig. 1G-H showed that tumor cDC1s expressed marked levels of *Xbp1s,* which was even superior to the levels noticed in bone-marrow derived DCs (BMDC) stimulated with the pharmacological UPR inducer tunicamycin. Altogether, these data indicate that cDC1s display prominent activation of the IRE1/Xbp1s axis in tumors.

We next interrogated whether the augmented IRE1 RNase activity observed in tumor cDC1s is a lineage-intrinsic signature or if it is a feature imposed by the tumor microenvironment. We quantified the VenusFP MFI of cDC1s directly exposed to the tumor (tumor cDCl and migratory cDC1s of the tumor draining lymph node (TdLN)), versus TdLN resident cDC1s, which do not access to the tumor site (gating strategy in Supp. Fig. 2B) (35). Data depicted in Fig. 1I indicates that tumor cDC1s express lower levels of VenusFP than resident cDC1s, indicating that tumor exposure does not increase IRE1 RNase activity in these cells. Similar observations were made for cDC2s (Fig 1I). In fact, cDC1s infiltrating MC38 tumors express markedly lower levels of VenusFP than spleen cDC1s (Supp. Fig. 1C). Of note, this observation was not replicated in monocytes, which express higher VenusFP levels in the tumor compared to the spleen (Fig 1J). Altogether, these data suggest that IRE1 RNase activation in tumor cDCs is a stable lineage intrinsic trait not driven by the microenvironment.

### XBP1 deletion in CD11c-expressing cells results in increased melanoma tumor growth

To gain insights on the role of XBP1 in DCs during melanoma tumor growth, we studied the *Itgax*-Cre x *Xbp1*^fl/fl^ mice (36, 37), referred to as ‘XBP1^ΔDC^’ mice, in which exon 2 of *Xbp1* is excised in CD11c-expressing cells, resulting in absence of the transcription factor in DCs (26). These animals were compared to control littermates (*Xbp1*^fl/fl^ animals with no expression of Cre), referred to as ‘XBP1^WT^’ mice. XBP1^WT^ and XBP1^ΔDC^ mice were implanted with the B78-ChOVA melanoma line, a B16 variant that expresses the ovalbumin (OVA) antigen and mCherry fluorescent protein (2). Data in Figure 2A-B indicate that XBP1^ΔDC^ mice showed moderate but noticeable acceleration of tumor growth and significantly larger tumor size than tumors from XBP1^WT^ mice on day 12 post implantation (80.44 ± 8.927 vs 56.56 ± 6.174 mm^3^, p = 0.0312, mean ± s.e.m.) (Fig. 2B). As a second tumor model, we also analyzed growth of subcutaneous MC38 murine colon adenocarcinoma tumors, which also showed a trend towards increased tumor growth in XBP1^ΔDC^ mice, but without reaching statistical significance (Supp. Fig. 3A-C).

**Figure 2.**
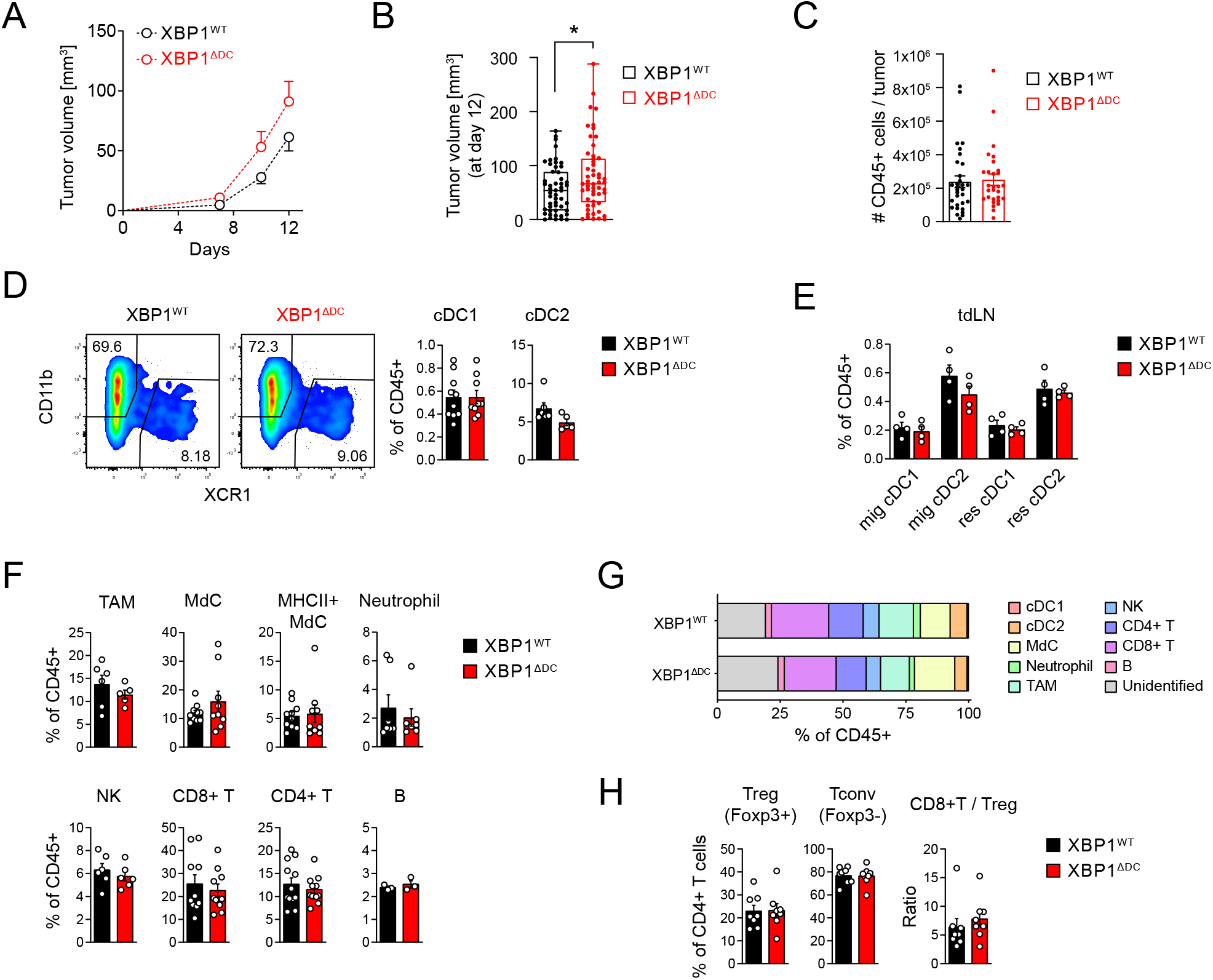
XBP1 deletion in CD11c-expressing cells results in increased melanoma tumor growth. XBP1^WT^ and XBP1^ΔDC^ mice were implanted with B78ChOVA tumors. **(A)** Tumor growth curves monitored over a period of 12 days. n=19 mice per group. Pooled data from 4 independent experiments. **(B)** Tumor size at day 12 post implantation. * p<0.05, two-tailed t-test. n= 51 mice (XBP1^WT^) or 53 mice (XBP1^ΔDC^) from animals used throughout this study (pooled data from 12 independent experiments), boxplot. **(C)** Cell counts for tumor immune infiltrate (CD45+). n=31 mice (XBP1^WT^) or 29 mice (XBP1^ΔDC^), pooled data from 8 independent experiments, mean ± s.e.m. **(D)** Frequencies of intratumoral cDC subsets. n=5-10, pooled data from two (cDC2) or three (cDC1) independent experiments, mean ± s.e.m. (E) Frequencies of cDC subsets in TdLN. n=4 mice per group, representative of two independent experiments, mean ± s.e.m. **(F-G)** Frequencies of tumor lymphoid and myeloid populations. n=3-12 mice per group, pooled data from two (neutrophils, TAMs, NK) or three (MdCs, T) independent experiments, mean ± s.e.m. **(H)** Tumor Treg (CD3^+^CD4^+^Foxp3^+^), Tconv (CD3^+^CD4^+^Foxp3^-^) and CD8^+^ T cells (CD3^+^CD8^+^) frequencies. n=8 mice per group, pooled data from two independent experiments, mean ± s.e.m.

To understand if XBP1 deficiency in the CD11c compartment resulted in altered cell recruitment, we quantified the immune cell composition at the melanoma tumor site. XBP1^WT^ and XBP1^ΔDC^ mice show similar numbers of CD45^+^ cells (Fig. 2C), and comparable composition of tumor cDC1/cDC2 and resident and migratory cDC1/cDC2s in the TdLN (Fig. 2D-E). Furthermore, conditional *knock-out* and control animals also showed comparable frequencies of myeloid and lymphoid cells (Fig. 2F-H). These data indicate that XBP1s expression by DCs infiltrating melanoma and MC38 tumor does not promote tumor progression. Rather, loss of XBP1s in DCs leads to increased melanoma tumor growth by mechanisms that are independent of immune cell recruitment.

### XBP1 deletion in CD11c-expressing cells results in impaired antitumor T cell responses and disbalanced precursor/terminal exhausted T cell ratio

We next focused our analysis on DC function. It was recently identified a conserved immunoregulatory transcriptional program activated by tumor DCs based on the co-expression of the molecules CD40 and PDL-1 plus the cytokine IL-12 (15). Our analysis revealed that tumor and migratory cDC1/cDC2s from XBP1^ΔDC^ mice express normal levels of these molecules (Fig. 3A, Supp. Fig. 4A). We also studied bone marrow cDC1s generated upon culture with OP9-DL1 stromal cell line plus FLT3L (reported in (38, 39)). We found that bone marrow cDC1s from XBP1^ΔDC^ mice produce lower levels of IL-l2 in unstimulated conditions (Fig 3B, Supp. Fig. 4B). However, upon tumor exposure, expression of IL-12 was comparable between XBP1 sufficient and deficient cDC1s. We conclude that Xbp1s regulate certain parameters of DC activation in steady state that are restored upon tumor encounter.

**Figure 3.**
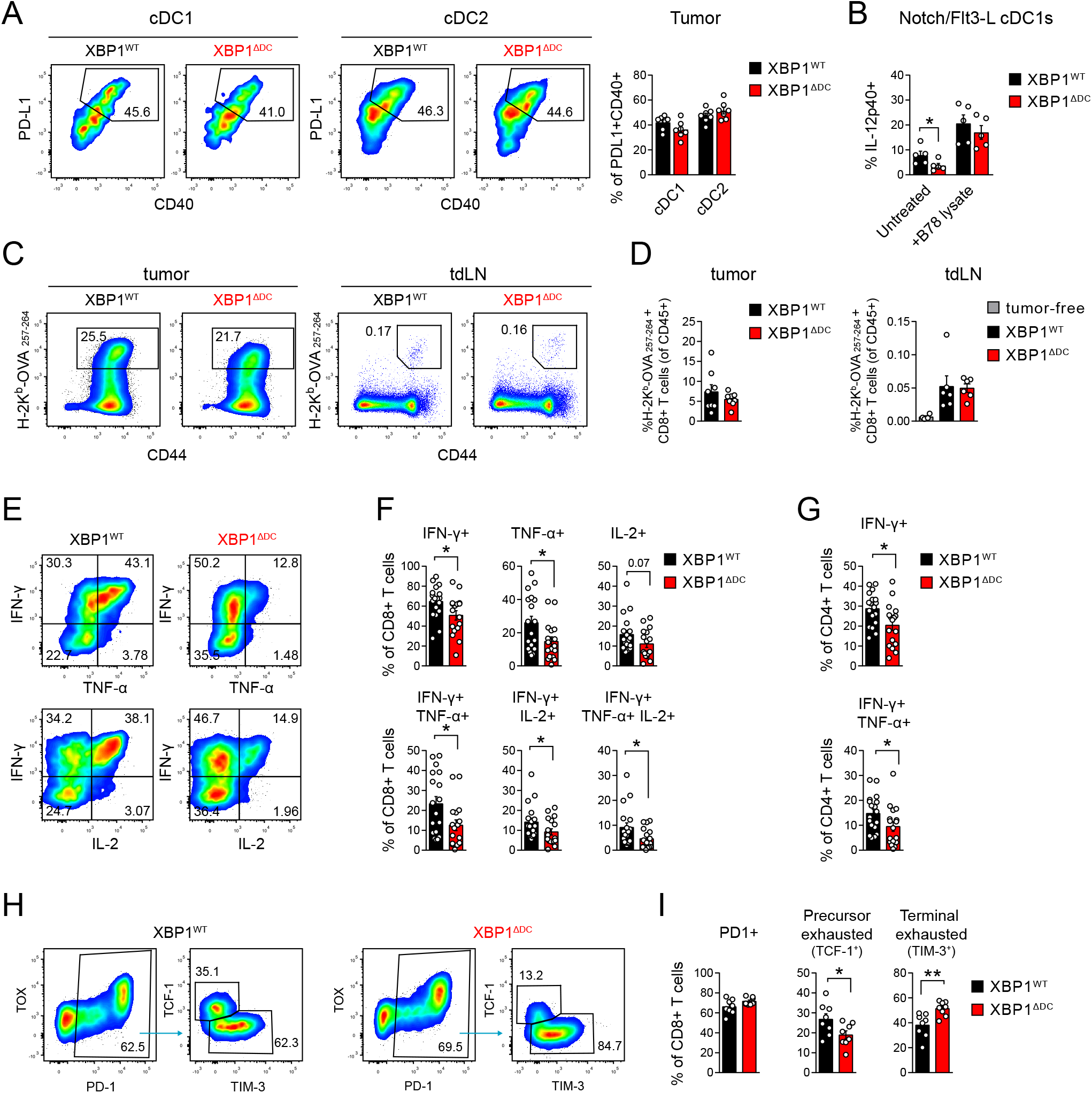
XBP1 deletion in CD11c-expressing cells results in impaired antitumor T cell responses and dysbalanced precursor/terminal exhausted T cell ratio. **(A)** CD40 and PD-Ll expression by tumor cDCs from B78-ChOVA bearing XBP1^WT^ or XBP1^ΔDC^ mice. n=7 mice per group, pooled data from two independent experiments, mean ± s.e.m. See also TdLN data in Supp. Fig 4A. **(B)** Intracellular IL-12 expression of *in vitro* generated cDC1s (FLT3-L/OP9-DL1) upon stimulation with B78-ChOVA lysates. Gated on cDC1s (MHC-II^+^ CD11c^+^ CD8^+^ CD11b^-^). *p < 0.05, two-tailed Mann-Whitney test. Each dot represents a biological replicate, n=5, data pooled from three independent experiments, mean ± s.e.m. (**C-D**) Tetramer^+^ CD8+ T cell frequencies in tumor and TdLN from B78-ChOVA bearing XBP1^WT^ or XBP1^ΔDC^ mice. Gated on CD3^+^CD8^+^ T cells. n=8 mice per group, pooled data from two independent experiments, mean ± s.e.m. **(E-G)** Cytokines expression by tumor CD8^+^ T cells (E-F) or CD4^+^ T cells (G) from B78-ChOVA bearing XBP1^WT^ or XBP1^ΔDC^ mice after ex vivo stimulation with PMA/ION in the presence of BFA. *p < 0.05, two-tailed Mann-Whitney test. n=18 mice (XBP1^WT^) or 17 mice (XBP1^ΔDC^), pooled data from 4 independent experiments, mean ± s.e.m. **(H-I)** Precursor exhausted (PD1^+^TCF1^+^ TIM3^neg^) and terminal exhausted (PD1^+^TCFl^neg^ TIM3^+^) CD8^+^ T cell tumor frequencies from B78-ChOVA bearing XBP1^WT^ or XBP1^ΔDC^ mice. Gated on CD3^+^CD8^+^ T cells. * p<0.05, **p<0.01, two-tailed t-test. N=8 mice per group, pooled data from two independent experiments, mean ± s.e.m.

Considering previous findings showing that Xbp1s deficient cDC1s have impaired cross-presentation abilities in steady state (26), we interrogated if the mice line was able to cross-present melanoma-associated antigens. We implanted B78ChOVA tumors in XBP1^WT^ and XBP1^ΔDC^ mice and quantified the presence of endogenous OVA-specific CD8^+^T cells using H-2K^b^-OVA_257-264_ tetramers. XBP1^WT^ and XBP1^ΔDC^ contained similar frequencies of OVA-specific CD8^+^ T cells in tumors and TdLN (Fig. 3C-D, Supp. Fig 4C). A similar response was obtained when tracking proliferation/early activation of CD8^+^ T cells isolated from pmel mice, which possess transgenic CD8^+^ T cells bearing a TCR selective for the melanoma-associated antigen gp100 (40) (Supp. Fig. 4D-E). Thus, we conclude that XBP1 deletion in tumor-associated CDllc^+^ cells does not prevent cross-presentation of melanoma-associated antigens.

We next investigated the quality of the antitumor T cell response evoked in XBP1^ΔDC^ mice. As a measure of T cell quality, we analyzed cytokine producing T cells from tumors of XBP1^WT^ and XBP1^ΔDC^ mice. Tumors from XBP1^ΔDC^ mice contained lower frequencies of IFN-γ-producing and TNF-producing CD8^+^ T cells, which also resulted in decreased frequencies of double producers IFN-γ^+^TNF^+^ CD8^+^ T cells and triple producers IFN-γ^+^TNF^+^IL-2^+^ CD8^+^ T cells (Fig. 3E-F). These observations were also noticed in the CD4^+^ T cell compartment, as reduced frequencies of IFN-γ^+^ CD4^+^ T cells and IFN-γ^+^ TNF^+^ CD4^+^ T cells were found in tumors from XBP1^ΔDC^ mice (Fig. 3G). As such, absence of XBP1 in CD11c-expressing cells results in decreased CD8^+^ and CD4^+^ T cell effector function in melanoma tumors. Analysis of the MC38 model showed that whereas the CD8^+^ T cell response was not affected, there was a significant reduction in the frequencies of IFN-γ-producing and IFN-γ/ TNF-producing CD4^+^ T cells (Supp. Fig. 4F-G).

Impaired cytokine production is a hallmark of CD8^+^ T cell exhaustion in cancer (41, 42), a process characterized by a progressive loss of function that culminates with the generation of terminal exhausted TIM-3^+^CD8^+^ T cells unable to control tumor growth (42). TIM-3^+^CD8^+^ T cells do not proliferate, are unresponsive to anti-PD1 therapy (41, 43) and originate from ‘precursor exhausted’ CD8^+^ T cells, a T cell state characterized by the expression of the transcription factor TCF-1 (termed ‘TCF1^+^CD8^+^ T cells’), which retain proliferative potential and can be reinvigorated through anti-PD1 therapy (41, 43, 44). We determined the presence of intratumoral TCF-1^+^CD8^+^ T cells and TIM-3^+^CD8^+^ T cells in melanoma tumors from XBP1^ΔDC^ mice and control animals. Tumors from XBP1^ΔDC^ mice show decreased infiltration of TCF-1^+^CD8^+^ T cells and increased proportions of TIM-3^+^CD8^+^ T cells compared to tumors from control animals (Figure 3H-I). Additionally, TIM3^+^ CD8^+^ T cells from XBP1^WT^ and XBP1^ΔDC^ mice display a *bona-fide* terminal exhausted phenotype, with elevated levels of CD39, TOX and granzyme B (Supp. Fig. 4H). These findings are consistent with data depicted in figure 3E-F showing lower frequencies of polyfunctional cytokine-producing CD8^+^ T cells in tumors from XBP1^ΔDC^ mice, which is an attribute of precursor exhausted TCF-1^+^CD8^+^ T cells (41). Altogether, our data shows that XBP1s in the CD11c^+^ compartment coordinates the balance of CD8^+^ T cell profiles in melanoma.

### Tumor cDC1s from XBP1^ΔDC^ mice display signs of RIDD

Thus far, our data indicate that XBP1s expression in DCs modulates melanoma tumor growth and the balance of effector and exhausted T cell subsets. However, it is unclear if these processes depend on XBP1s-transcriptional activity, as previous work demonstrated that XBP1s deficiency leads to hyperactivation of IRE1 RNase and RIDD in steady state cDC1s (25, 26). Thus, we interrogated if XBP1 deficiency also resulted in increased IRE1 RNase activity in tumor cDC1s (Fig 4A). We measured expression of *Xbp1* spliced/unspliced mRNA in tumor cDC1s from control and XBP1^ΔDC^ mice. Although XBP1^ΔDC^ cDC1s are unable to synthesize XBP1s protein, these cells still generate *Xbp1* mRNA bearing the IRE1 cleavage sites, which serves as an assay to monitor IRE1 RNase activity (26). Data in Figure 4B show that tumor cDC1s isolated from XBP1^ΔDC^ mice express marked levels of *Xbp1s* mRNA compared to control counterparts, which is an indicative sign of IRE1 RNase hyperactivation.

**Figure 4.**
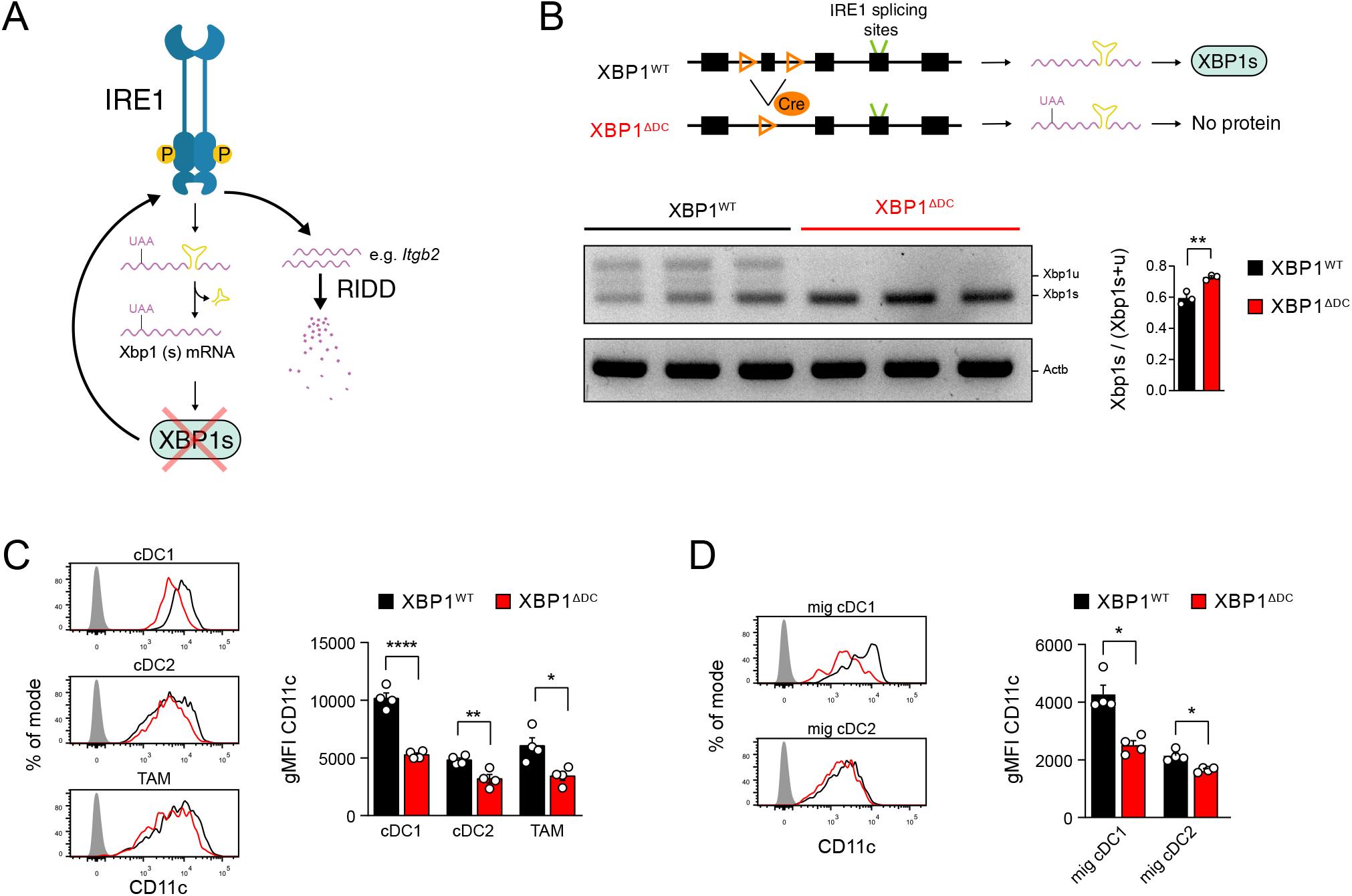
Tumor cDC1s from XBP1ΔDC mice display signs of RIDD. (A) Upon Cre mediated recombination in XBP1^fl/fl^ mice, a premature stop codon is introduced in the *XBP1* mRNA sequence, preventing the translation of a functional Xbp1s protein. XBP1s absence is reported to trigger IRE1 RNase hyperactivation and induce RIDD in certain cell types (26, 36). However, IRE1 RNase activity can still be monitored by determining Xbp1 mRNA splicing ratio. (B) Scheme depicting LoxP sites and IRE1 splicing sites at *XBP1* locus (top) and PCR analysis of Xbp1 splicing in intratumoral cDC1s isolated from B16-bearing XBP1^WT^ and XBP1^ΔDC^ mice (bottom). Each lane represents different mice. Xbp1u: Xbp1 unspliced; Xbp1s: Xbp1 spliced; Actb: beta actin. (C) CD11c expression by intratumoral cDC1, cDC2 and TAMs from B16-bearing XBP1^WT^ and XBP1^ΔDC^ mice. Gray histogram depicts unstained control. * p<0.05, ** p<0.01, **** p<0.0001, two-tailed t-test. n=4 mice per group, representative of two independent experiments, mean ± s.e.m. (D) CD11c expression by cDC subsets in the TdLN from B16-bearing XBP1^WT^ and XBP1^ΔDC^ mice. Gray histogram depicts unstained control. *p<0.05, two-tailed Mann-Whitney test. n=4 mice per group, representative of two independent experiments, mean ± s.e.m.

To determine RIDD on protein level in tumor DCs, we determined surface expression of the integrin CD11c, a dimeric partner of *Itgb2* (coding the integrin CD18), which is a reported mRNA substrate of IRE1 RNase (26). CD11c surface expression depends on RIDD-mediated degradation of *Itgb2* mRNA and therefore, it can be used as a surrogate marker for RIDD activity. Data in Fig. 4C show that tumor cDC1s from XBP1^ΔDC^ express lower surface levels of CD11c than control counterparts, confirming RIDD induction on protein level. Similar effect was observed in cDCl subsets from the TdLN (Fig 4D). These data show that XBP1-deficient cDC1s display signs of RIDD in melanoma tumors. Interestingly, additional APCs such as tumor cDC2s and TAM from XBP1^ΔDC^ mice showed a modest but noticeable reduction in CD11c expression (Fig. 4C), suggesting that these cells may also induce RIDD upon XBP1 loss, albeit at lower extent than cDC1s.

### Gene expression profiles of tumor cDC1s deficient for IRE1 RNase and XBP1

Given that XBP1 deficient cDC1s show signs of RIDD at the tumor site, we analyzed the transcriptomic signature downstream of IRE1 RNase in melanoma-infiltrating cDC1s. To identify specific XBP1-dependent and RIDD-dependent targets, we carried out a parallel analysis of the transcriptome of tumor cDC1s deficient for XBP1, or double deficient for the RNase domain of IRE1 and XBP1. To generate double *knock-out* animals for the IRE1 RNase and XBP1 in CD11c-expressing cells, we crossed XBP1^ΔDC^ mice with *Ern1*^fl/fl^ mice, which bear loxP sites flanking exon 20 and 21 of the *Ern1* gene and generates a truncated IRE1 isoform lacking the RNase domain (45) (referred to as “XBP1^ΔDC^/IRE1^trunc^DC mice”). As such, XBP1^ΔDC^ mice lack the transcription factor and activate RIDD, whereas double deficient XBP1^ΔDC^/IRE1^trunc^DC mice lack both Xbp1s and RIDD (Supp. Fig 5A-C). With this strategy, Xbp1s target genes are identified as transcripts that are downregulated in tumor cDC1s from both XBP1^ΔDC^ and XBP1^ΔDC^/IRE1^trunc^DC mice. In contrast, RIDD-dependent transcripts are recognized as mRNAs that decrease their expression in XBP1-deficient tumor cDC1s, but which expression is restored in XBP1^ΔDC^/IRE1^trunc^DC animals.

Tumor cDC1s were isolated by cell sorting from B16 melanoma tumors of control, XBP1^ΔDC^ or XBP1^ΔDC^/IRE1^trunc^DC mice and the transcriptome was analyzed by bulk RNA sequencing (RNA-seq). 51 differentially expressed genes (DEG) were identified among XBP1^ΔDC^ or XBP1^ΔDC^/IRE1^trunc^DC cDC1s (adjusted p-value < 0.05 and |Fold Change| > 1.5) (Fig. 5A). Biological pathway enrichment analysis using Gene Ontology (GO) knowledgebase revealed that a large proportion of DEGs are constituents of the response to misfolded proteins and transport and localization of ER proteins (see Biological Process, Fig. 5B). There was also an overrepresentation of protein disulfide-isomerases (see Molecular Function, Fig. 5B) and on intracellular localization level, most deregulated genes encoded ER proteins (see Cellular Component, Fig. 5B). Next, we analyzed the DEGs per mice line, which clustered these genes into three main groups: (1) genes upregulated in XBP1-deficient and IRE1 RNase/XBP1-deficient tumor cDC1s; (2) genes downregulated in both XBP1-deficient and IRE1 RNase/XBP1-deficient tumor cDC1s (XBP1s targets); and (3) genes downregulated exclusively in XBP1-deficient tumor cDC1s (potential RIDD targets) (Fig. 5A). In group (1), we identified five transcripts including *Hspa5* which encodes BiP, a chaperone induced upon UPR activation (46). This finding indicates that XBP1-deficient and IRE1 RNase/XBP1-double deficient tumor cDC1s show signs of ER stress (Fig. 5A). We also identified *Cox6a2* (subunit of cytochrome C oxidase), which was previously identified as an upregulated gene in XBP1-deficient cDC1s (26). Transcripts in group (2) include protein disulfide isomerases (*Erp44, Txndx11, Txndc5, P4hb*) chaperones (*Dnajb9/Hyou1*); glycosylation proteins (*Rpn1, Alg2, Serp1*), proteins involved in transport to the ER (*Sec61a1, Sec61b, Spcs2, Spcs3, Ssr3*) and from the ER to Golgi (*Bet1, Surf4*) (Fig. 5A). Additional canonical XBP1s targets (*Stt3a* and *Edem2*) were identified when the cut off value was set below 1.5-fold (Supp. Fig 6A). Gene Set Enrichment Analysis (GSEA) revealed that the transcriptome of both XBP1^ΔDC^ and XBP1^ΔDC^/IRE1^trunc^DC tumor cDC1s were depleted of targets related to protein glycosylation, ER to Golgi transport, protein localization to the ER and lipid biosynthesis (Fig 5C-D).

**Figure 5.**
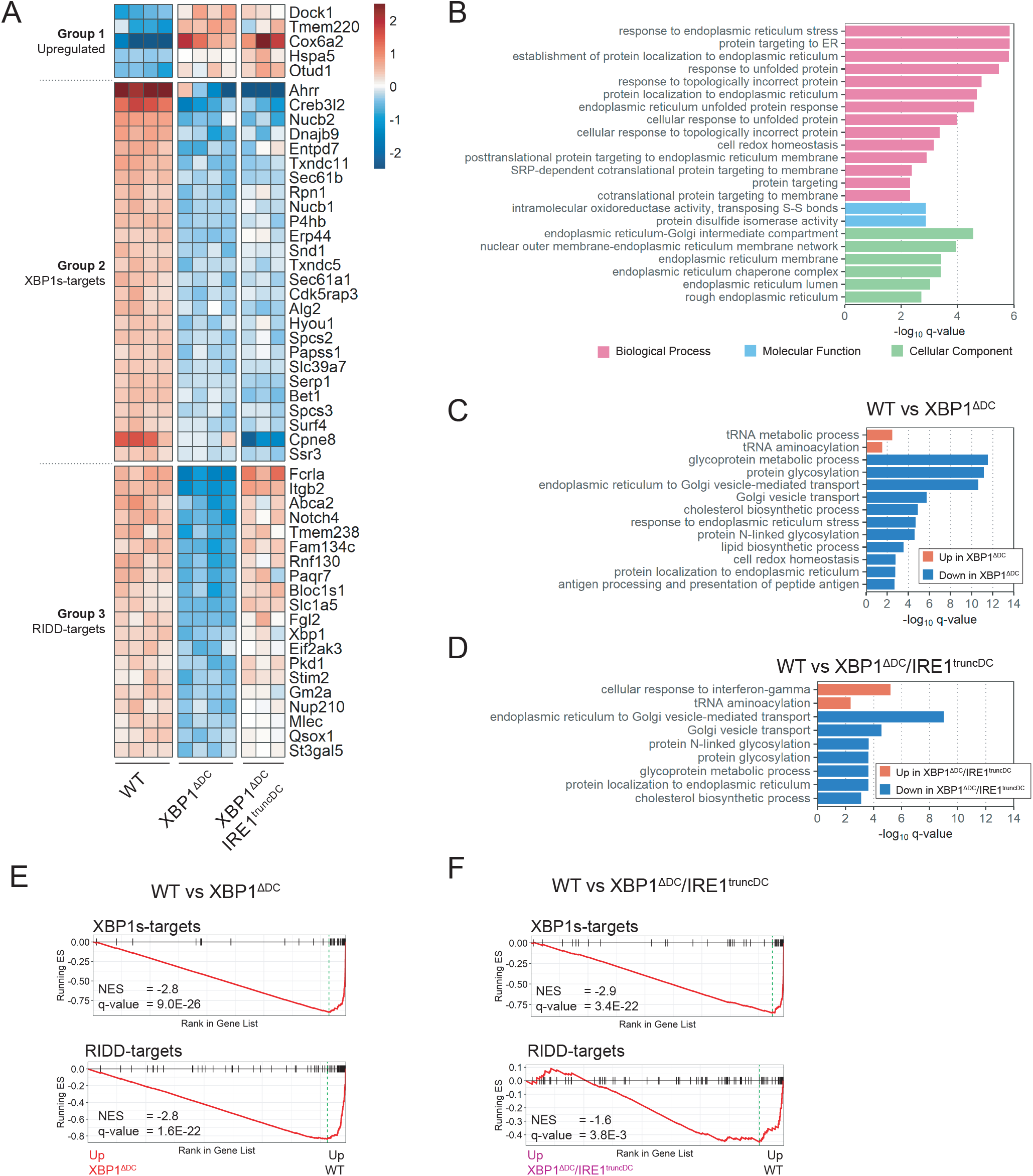
Gene expression profiles of tumor cDC1s deficient for IRE1 RNase and XBP1. WT, XBP1^ΔDC^ and XBP1^ΔDC^/IRE1^trunc^DC mice were implanted with B16 tumors. After 12 days, tumor cDC1s were isolated by cell sorting and total RNA was sequenced by RNA-seq. **(A)** Heatmap of differentially expressed genes (DEGs). Three groups of genes were identified by the pattern of expression among the three genotypes. **(B)** Over representation analysis of DEGs over the Gene Ontology (GO) database. **(C)** Gene Set Enrichment Analysis (GSEA) of WT vs XBP1^ΔDC^ cDC1s using GO:Biological Process database. **(D)** GSEA of WT vs XBP1^ΔDC^/IRE1^trunc^DC cDC1s using GO:Biological Process database. **(E-F)** GSEA using XBP1s- and RIDD-target gene sets from literature (So et al, 2012, Cell Metabolism).

Finally, group (3) includes the canonical RIDD substrates *Bloc1s1, St3ga15* (21), *Itgb2* (26), plus a subset of transcripts with heterogeneous functions (Fig. 5A) that range from lipid synthesis and metabolism members (*St3ga15, Gm2a, Abca2*), Ca^+2^ homeostasis (*Stim2, Pkd1*), protein folding (*Qsox1, Mlec*), a steroid binding receptor (*Paqr7*), an amino acid transporter (*Slc1a5*), a member of the nuclear pore complex (*Nup210*), an E3-ubiquitin ligase (*Rnf130*), signaling receptors in immunity and development (*Fcrla, Notch4*, respectively) to *Eif2ak3*, which encodes the UPR transducer PERK. Consistent with the functional heterogeneity of RIDD targets, GSEA did not reveal differences at the level of biological processes between XBP1^ΔDC^ and XBP1^ΔDC^/IRE1^trunc^DC cDC1s (Fig 5C-D). Even though some processes such as antigen processing and presentation and cell-redox homeostasis are downregulated exclusively in tumor cDC1s from XBP1^ΔDC^ mice (thereby suggesting RIDD dependency), these processes display low enrichment scores. To sum up, these data indicate that tumor cDC1s from XBP1^ΔDC^ mice display an altered XBP1s transcriptional program related to protein homeostasis and folding, and counter activate RIDD. Using GSEA and reported XBP1s-target and RIDD-target gene datasets (31, 47) we confirmed downregulation of the canonical XBP1s transcription program in both XBP1^ΔDC^cDC1s and double deficient XBP1^ΔDC^/IRE1^trunc^DC cDC1s (Fig 5E-F, Supp. Fig. S6B), whereas RIDD-dependent targets are predominantly downregulated in XBP1^ΔDC^ cDC1s (Fig 5E-F, Supp Fig. S6B). Finally, our findings reveal that melanoma-infiltrating cDC1s do not show signs of dysfunctional XBP1s activity, as genes related to triglyceride biosynthesis that are associated with diminished DC function in other cancer settings are not downregulated upon XBP1s or IRE1 RNase loss (Supp. Fig. 6C-D) (30). Taken together, these results demonstrate that XBP1 deficiency in tumor cDC1s impairs transcriptomic programs associated with the maintenance of proteostasis and induces RIDD.

### RIDD activation in DCs accounts for the changes related to tumor growth and dysregulated antitumor T cell immunity noticed in XBP1^ΔDC^ mice

Our observations raise the question as to whether the increased tumor growth and T cell dysregulation noticed in XBP1^ΔDC^ mice is due to Xbp1s- or RIDD-dependent outputs in DCs. To address this issue, we implanted the B78ChOVA cell line in control and XBP1^ΔDC^/IRE1^trunc^DC animals (Fig. 6A-B), in which RIDD is abolished (Supp. Fig. 5A-C). Remarkably, in contrast to the observations noticed in XBP1^ΔDC^ mice (Fig. 2A), the tumor size of animals lacking both XBP1 and IRE1 RNase in DCs was comparable to that observed in control animals (41.78 ± 9.033 vs 53.90 ± 12.36 mm^3^, p=0.42, mean ± s.e.m.) (Fig. 6B), indicating that RIDD activation in CD11c-expressing cells accounts for the increased tumor growth. Furthermore, we examined whether the reduced frequencies of cytokine producing CD8^+^ T cells and the disbalance between TCF-1^+^/TIM-3^+^ CD8^+^ T cells noticed in XBP1^ΔDC^ mice are also dependent on RIDD. Analysis of tumor-infiltrating CD8^+^T cells revealed no differences in the frequencies of IFN-γ producing CD8^+^T cells, TNF producing CD8^+^T cells, IL-2 producing CD8^+^T cells, nor in the proportion of double or triple cytokine producers between XBP1^ΔDC^/IRE1^trunc^DC mice and control animals (Fig. 6C-D). Similar results were observed for IFN-γ producing or double IFN-γ/ TNF-α producing CD4^+^ T cells (Fig. 6E). In addition, analysis of the composition of precursor-exhausted/terminal exhausted T cells revealed similar infiltration of TCF-1^+^/TIM-3^+^ CD8^+^T cells in melanoma tumors from XBP1^ΔDC^/IRE1^trunc^DC mice versus control animals (Fig. 6F-G), indicating that IRE1 RNase activity in DCs accounted for the accumulation of dysfunctional CD8^+^ T cells in melanoma. Altogether, these data indicates that hyperactivation of the RNase domain of IRE1 in DCs fine tunes melanoma tumor growth and antitumor T cell immunity.

**Figure 6.**
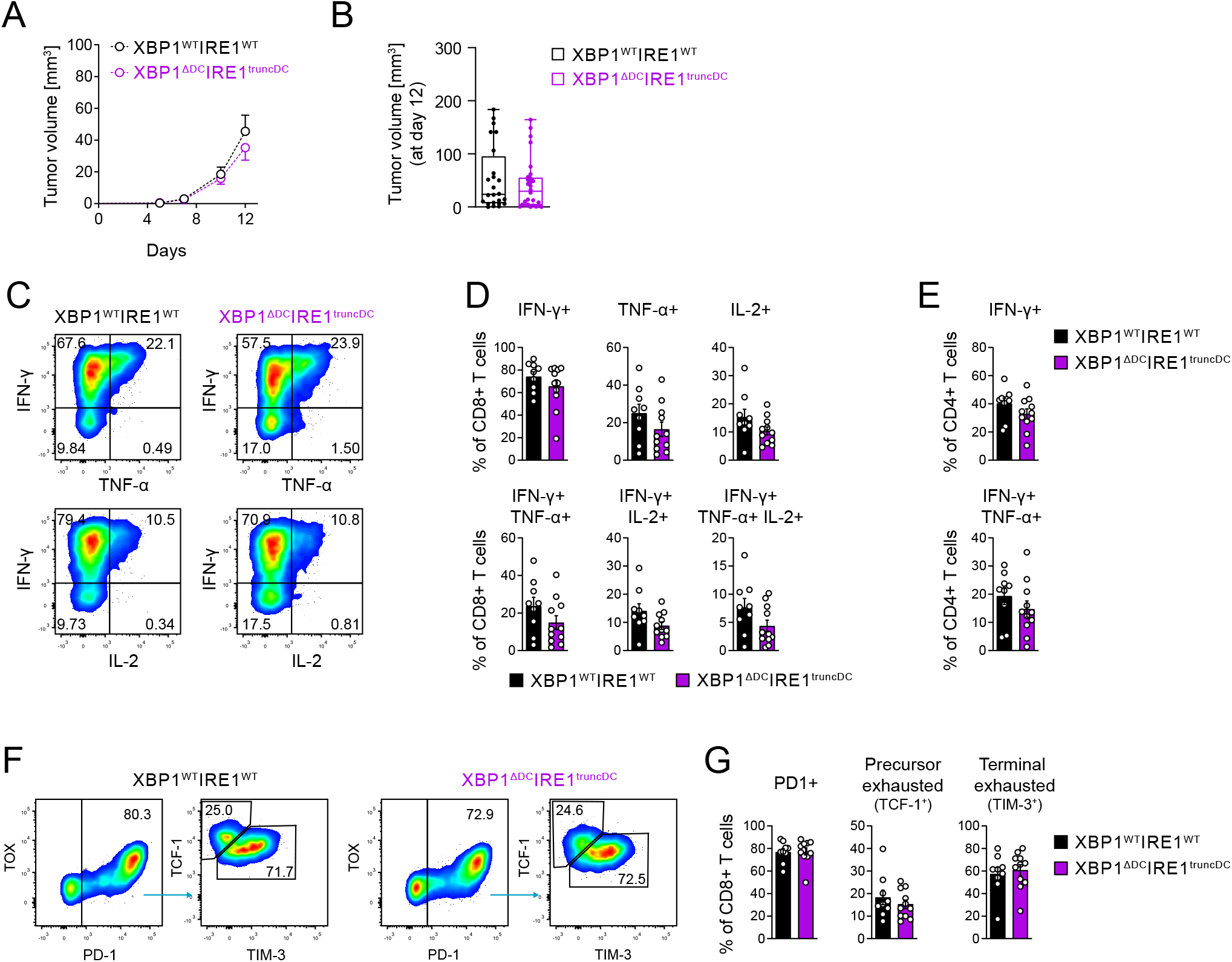
RIDD activation in DCs accounts for the changes related to tumor growth and dysregulated antitumor T cell immunity noticed in XBP1ΔDC mice. XBP1^WT^/IRE1^WT^ and XBP1^ΔDC^/IRE1^trunc^DC mice were implanted with B78ChOVA cells. **(A)** Tumor growth curves monitored over a period of 12 days. n=20 mice (XBP1^WT^/IRE1^WT^) or 24 mice (XBP1^ΔDC^/IRE1^trunc^DC), data pooled from 4 independent experiments. **(B)** Tumor size at day 12 post implantation. n=24 mice (XBP1^WT^/IRE1^WT^) or 28 mice (XBP1^ΔDC^/IRE1^trunc^DC), data pooled from 5 independent experiments, mean ± s.e.m. **(C-D)** Cytokines expression by tumor CD8^+^ T cells after ex vivo stimulation with PMA/Ionomycin in the presence of BFA. Gated on CD3^+^CD8^+^ T cells. N=9 mice (XBP1^WT^/IRE1^WT^) or 11 mice (XBP1^ΔDC^/IRE1^trunc^DC), data pooled from three independent experiments, mean ± s.e.m. **(E-F)** Precursor exhausted (PD1^+^TCF1^+^TIM3^neg^) and terminal exhausted (PD1^+^TCF1^neg^ TIM3^+^) CD8^+^ T cell tumor frequencies. Gated on CD3^+^CD8^+^ T cells. n=9 mice (XBP1^WT^/IRE1^WT^) or 11 mice (XBP1^ΔDC^/IRE1^trunc^DC), data pooled from three independent experiments, mean ± s.e.m.

## DISCUSSION

The IRE1/XBP1s axis has emerged as a critical regulator of immunity and cancer (27, 48, 49). The differential mechanisms by which IRE1 signaling integrates the intensity and duration of ER stress to regulate cell fate is particularly noticed in the immune system, with cells such as cDC1s, B cells, NK cells and eosinophils that opt for an intact IRE1/XBP1s axis to maintain cellular health (25, 31, 50, 51), or cells including TAM/MdCs or intratumoral T cells, which acquire dysfunctional phenotypes upon enforced activation of the UPR sensor (27, 32, 52).

Here, we report that loss of XBP1 in CD11c-expressing cells results in increased melanoma tumor growth, decreased frequencies of cytokine-producing T cells and accumulation of terminal exhausted TIM-3^+^CD8^+^ T cells. Notably, this effect is abrogated in XBP1^ΔDC^/IRE1^trunc^DC mice, demonstrating that IRE1 RNase-dependent, XBP1s-independent outputs account for the dysregulated antitumor immunity in melanoma. We also observe a milder phenotype in the MC38 model, suggesting that different tumor models differentially regulate the IRE1/XBP1s axis in DCs.

The results presented in this work contrast with previous studies demonstrating that persistent IRE1/XBP1s activation in tumor DCs curtails their antitumor function. In ovarian cancer, the same XBP1^ΔDC^ mice line show marked inhibition of tumor progression and improved antitumor immunity (30). One possibility accounting for these differences may be related to the different composition of DCs infiltrating these cancer models. In ovarian cancer models, tumor-associated DCs are spontaneously immunosuppressive (53) and in fact, whereas depletion of CD11c^+^ cells delays ovarian cancer progression in later stages (54), the same process curtails CD8^+^ T cell priming in melanoma (55). Furthermore, XBP1 deficiency in ovarian cancer DCs does not lead to RIDD activation (30), which contrasts to the evidence presented in this work. Thus, as result of these combined data, we must consider that the outcome of the IRE1 outputs in tumor DCs may drastically differ depending on the DC subset and the cancer type. In fact, data presented here show that animals lacking both XBP1 and IRE1 RNase in DCs display normal melanoma tumor growth and T cell immunity, indicating that the deletion of an entire branch of the UPR in DCs does not have a predetermined role across different tumor types. These results complement existing literature and may be relevant for building a comprehensive understanding on future manipulation of the IRE1/XBP1s axis in cancer. In addition, even though we use a genetic model of XBP1 deletion to reveal the scope of RIDD in tumor DCs which may not recapitulate physiological responses, our work alludes to cDC1s as regulators of anti-melanoma T cell immunity in XBP1^ΔDC^ mice. First, cDC1s are the subset with the highest IRE1 RNase activity within melanoma and MC38 tumor niches, and T cell parameters altered by XBP1 loss in CD11c^+^ cells, such as the induction of intratumoral IFN-γ producing CD8^+^ T cells and the maintenance of precursor exhausted TCF1^+^CD8^+^ T cells are attributed to cDC1 function (8, 11). Furthermore, these processes are also dependent on RIDD activation by DCs, which is more strongly induced in cDC1s from XBP1^ΔDC^ mice (25, 26). Nevertheless, as we also observe reduced frequencies of tumor infiltrating CD4^+^ IFNγ^+^ T cells in XBP1^ΔDC^ mice in melanoma and MC38 models, a contribution of IRE1 RNase in CD4^+^ T cell priming by tumor cDC1s (56) or also by cDC2s (17) cannot be excluded.

On a mechanistic level, we do not find a role for IRE1 RNase in cross-presentation of tumor antigens, contrasting with previous findings in steady state cDC1s (26). These data suggest that tumors may shape the spectra of XBP1s/RIDD targets in infiltrating cDC1s or that additional mechanisms (or DC subtypes) may compensate for the process. In fact, *tapbp* mRNA, a previously identified RIDD target in XBP1-deficient splenic DCs contributing to antigen cross-presentation is not found as DEG in the transcriptomic analysis of this study. In addition, growing evidence demonstrating the capacity of tumor DCs to carry out cross-dressing of MHC-I/peptide complexes from tumor cells adds a new layer of complexity that remains to be addressed (57, 58). However, one novel candidate identified in this analysis as a potential RIDD substrate, the ER-resident FC receptor Like A (*Fcr1a*) (Fig. 5A), has been previously identified as part of a BATF3/IRF8 transcriptional program that confers tumor immunogenicity in cDC1s independently of cross-presentation (59). As such, data presented here delineates for first time the XBP1s-dependent and RIDD-dependent targets in tumor cDC1s, which may serve as basis for future studies focused on addressing the role of selective IRE1 RNase targets involved in the regulation of antitumor immunity.

Multiple efforts are currently focused on the development of pharmacological compounds targeting the IRE1 RNase active site and XBP1s *in vivo*, many of which have shown translational potential in cancer (60). A recent study revealed that RIDD regulates expression of the MHC-I heavy chain mRNAs in DCs and that inhibition of IRE1 RNase through systemic administration of small molecules greatly attenuates tumor growth in 4Tl and CT26 models, by a mechanism proposed to be dependent on DC cross-presentation (61). Even though we do not find MHC-I heavy chain mRNAs as DEGs in the transcriptomic analysis of tumor cDC1s from XBP1^ΔDC^/IRE1^trunc^DC mice, and we do not find an improved melanoma tumor response in XBP1^ΔDC^/IRE1^trunc^DC animals, future studies are required to integrate these findings through experiments that include kinetics of comparable tumor models. Finally, the work presented here serves as a proof-of-concept study demonstrating that IRE1 RNase dependent, XBP1s-independent outputs in DCs may also contribute to fine-tuning antitumor immunity.

## MATERIAL AND METHODS

### RESOURCE AVAILABILITY

Further information and requests for resources and reagents should be directed to Fabiola Osorio.

#### Materials availability

This study did not generate new unique reagents.

#### Data and code availability

RNA-seq data have been deposited at GEO and are publicly available as of the date of publication. Accession numbers are listed in supplementary resources table. This paper does not report original code. Any additional information required to reanalyze the data reported in this paper is available upon request.

### EXPERIMENTAL MODEL AND SUBJECT DETAILS

#### Mice

ERAI (33), XBP1^WT^(XBP1fl/fl (36)), XBP1^ΔDC^ (XBP1fl/fl x CD11c-Cre (37)), XBP1^WT^/IRE1^WT^ (XBP1fl/fl x IRE1fl/fl (45)), XBP1^ΔDC^ IREU™°DC (XBP1fl/fl x IRE1fl/fl x CD11c-Cre) mice were bred at Universidad de Chile and Fundación Ciencia y Vida in specific pathogen-free conditions. Also, for RNA seq studies XBP1^WT^, XBP1^ΔDC^, XBP1^WT^/IRE1^WT^ and XBP1^ΔDC^ IRE1^trunc^DC mice were bred at the animal facility at VIB institute. pmel-1 mice (40) were kindly donated by Dr F. Salazar-Onfray. All mice were kept on a C57BL/6 background. Litters with mice of both sexes at 6–14 weeks of age were used for experiments.

#### Cell Lines

B78-ChOVA cells were kindly provided by Dr. Matthew Krummel (UCSF) (2). B16-F10 cells were obtained from ATCG (#CRL-6475). B16-FLT3L cell line (62) was provided by Dr. Maria Rosa Bono (University of Chile). MC-38 cell line was provided by Dr. Álvaro Lladser (Universidad San Sebastian). OP9 cells expressing Notch ligand DL1 (OP9-DL1) (63) were kindly provided by Dr. Juan Carlos Zuñiga-Pflucker (Sunnybrook Research Institute, Canada). Cells were cultured under standard conditions prior to injection into mice. Briefly, cells were cultured in DMEM (B78-ChOVA) or RPMI-1640 (B16-F10/B16-FLT3L/MC-38) supplemented with 10% v/v inactivated fetal bovine serum (FBS, Gibco), 100 U/mL penicillin (Corning), 100 μg/mL streptomycin (Corning) and 0.55 mM 2-Mercaptoethanol (Gibco). For MC-38 culture, media was supplemented additionally with non-essential amino acids (ThermoFisher Scientific) and 1 mM sodium pyruvate (ThermoFisher Scientific). Cells were cultured on T75 tissue-culture treated plastic flasks at 37°C, 5% CO_2_. Cells were split every other day. OP-DL1 cells were cultured in MEM-alpha medium supplemented with 20% FBS (Gibco), 100 U/mL penicillin (Corning), 100 μg/mL streptomycin (Corning), 1mM sodium pyruvate (brand) and 0.55 mM 2-Mercaptoethanol (Gibco).

### METHOD DETAILS

#### Tumor Model

Tumor cell lines were harvested, washed with PBS, and resuspended in a final injection volume of 50 μl PBS. 5×10^5^ (B16/B78-ChOVA) or 1×10^6^ (MC-38) tumor cells were injected in the right flank of shaved mice intradermally and allowed to grow for 10-15 days. For tumor growth curves, tumor size was determined by two orthogonal measurements with a caliper and the volume was estimated as (width^2 x length)/2.

#### Preparation of Cell Suspensions

Tumors were minced and digested with Collagenase D (1 mg/mL, Roche) and DNAse I (50 μg/mL, Roche) for 30 minutes at 37°C in a water bath. Digested tissue was then passed through a 70 μm cell strainer, followed by red blood cell lysis with RBC lysis buffer (Biolegend). Single cells were kept on ice.

For whole intratumoral immune cell profiling and DC stainings, CD45-biotin magnetic positive selection (MACS, Miltenyi) was performed to enrich for total tumor immune infiltrate.

For intratumoral T cell stainings, hematopoietic cells were enriched by density gradient centrifugation with 40/70 Percoll (GE Healthcare) for 20 min at 700xg.

Tumor draining lymph nodes (tdLNs) were minced and digested with Collagenase D (1 mg/mL, Roche) and DNAse I (50 μg/mL, Roche) for 45 minutes at 37°C in a water bath. Digested tissue was then passed through a 70 μm cell strainer and single cells were kept on ice.

#### Bone marrow derived DCs generation

Bone marrow cells from femurs and tibias were cultured in presence of 20 ng/ml mouse recombinant GM-CSF (Biolegend) for 8 days. Fresh culture medium with cytokine was added on day 3, and on day 6. After harvesting and when indicated, cells were stimulated with 1 ug/mL Tunicamycin (Sigma) for 8h followed by total RNA extraction with Trizol (Invitrogen).

#### Bone marrow derived cDC1s generation and tumor lysate stimulation

Bone marrow cells from femurs and tibias were cultured in presence of 100 ng/ml recombinant human FLT3-L (Peprotech). After three days of differentiation, cells were plated onto a monolayer of OP9-DL1 stromal cells and co-cultured for additional 6 days in P24 plates as previously reported (38, 39).

For tumor lysate preparation B78-ChOVA cells were washed twice with PBS, resuspended at 8×10^6^ cells/mL in RPMI supplemented with 10% FBS and aliquoted in cryotubes. Cell suspensions were subjected to heat-shock (42°C for 60 min) followed by three cycles of freeze/thaw (liquid nitrogen/waterbath at 37°C). Tumor lysates were stored at −80°C until use.

BM-derived cDC1s were harvested and plated with B78-ChOVA lysates (50 uL/mL) in round-bottom p96 plates. After 14h, Brefeldin A (GolgiPlug, BD) was added and four hours later, cells were harvested and stained for intracellular IL-12p40 expression by flow cytometry.

#### Xbp1s splicing assay

Total RNA was isolated either by Trizol (Invitrogen) or RNAeasy plus Micro Kit (Qiagen). cDNA was prepared using M-MLV reverse transcriptase (Invitrogen). The following primers were used for conventional PCR amplification of total Xbp1: Fwd: 5’-ACACGCTTGGGAATGGACAC-3’ and Rev: 5’-CCATGGGAAGATGTTCTGGG-3’ (26); and for beta actin (*Actb*): Fwd: 5’-CTAAGGCCAACCGTGAAAAG-3’ and Rev: 5’-TTGCTGATCCACATCTGCTG-3’ or alternatively for beta actin (*Actb*): Fwd 5’-GTGACGTTGACATCCGTAAAGA-3’ and Rev: 5’-GCCGGACTCATCGTACTCC-3’. PCR products were analyzed on agarose gels.

#### Flow Cytometry and Cell Sorting

For surface staining, cells were incubated with anti-Fc receptor antibody (anti-CD16/32, Biolegend) and then stained with fluorochrome-conjugated antibodies in FACS buffer (PBS + 1% FBS + 2mM EDTA) for 30 min at 4°C. Viability was assessed by staining with fixable viability Zombie (BioLegend) or LIVE/DEAD fixable (Invitrogen). A biotinylated antibody was used for F4/80 staining, followed by a second staining step with Streptoavidin-APC (Biolegend) for 30 min at 4°C. Flow cytometry was performed on BD Fortessa LSR instrument. Analysis of flow cytometry data was done using FlowJo software. Cell sorting was performed using a BD FACS Aria III.

#### Transcription Factors and Granzyme B intracellular staining

After surface staining, cells were fixed and permeabilized using Foxp3 transcription factor staining set (eBioscience) followed by intracellular staining of transcription factors (Foxp3, Tcf1, Tox) and/or granzyme B as indicated by the manufacturer protocol.

#### T cell stimulation and intracellular cytokine staining

Tumor and TdLN cell suspensions were stimulated *ex-vivo* prior to staining with 0.25 μM phorbol 12-myristate 13-acetate (PMA; Sigma) and 1 μg/mL Ionomycin (Sigma) at 37°C and 5% CO_2_ for 3.5 hr in the presence of Brefeldin A (BD GolgiPlug). After stimulation, cells were surface stained as mentioned above. Then, cells were fixed and permeabilized using BD Cytofix/Cytoperm fixation/permeabilization kit (BD) followed by intracellular staining of cytokines (IFN-γ, IL-2 and TNF-α) as indicated by the manufacturer protocol.

#### Tetramer staining

For OVA-specific CD8+ T cell quantification cells were incubated with PE H2-K^b^-OVA (SIINFEKL) tetramers (MBL) at room temperature for 30 min protected from light, followed by surface staining and FACS analysis.

#### t-SNE and clustering

For tSNE visualization of tumor immune infiltrate a multicolor flow cytometry panel was used including 19 parameters (FSC, SSC, Viability, CD45, VenusFP, XCR1, CD4, NK1.1, CD26, F4/80, Ly6G, MHCII, CD24, CD3e, Ly6C, CD8a, CD11c, CD11b, CD19). Cells were compensated for spillover between channels and pre-gated on CD45+ Live singlets using FlowJo. Flowjo workspace was imported into the R environment using CytoML v2.4.0, FlowWokspace v4.4.0 and FlowCore v2.4.0 packages (64–66). The intensity values of marker expression were then biexp-transformed via the flowjo_biexp_trans function of FlowWorkspace using parameters ChannelRange=4096, maxValue=262144, pos=4.5, neg=0 and widthBasis=-10. Subsequently 5.000 cell events from each mouse (4 WT and 4 ERAI) were randomly sampled and combined for a total of 40.000 single cells. Sampled data was min-max normalized, and subjected to dimensionality reduction by Barnes-Hutts implementation of t-Distributed Stochastic Neighbor Embedding (tSNE) using RtSNE v0.15 package (67). Thirteen parameters were used for tSNE construction (XCR1, CD4, NK1.1, CD26, F4/80, Ly6G, MHCII, CD24, CD3e, Ly6C, CD8a, CD11c, CD11b and CD19) and the parameters were set to iterations=1000 and perplexity =30. After dimensionality reduction, automatic clustering was performed using density based spatial clustering (DBSCAN) using DBSCAN v1.1.8 package (68). Dotplot for marker expression among clusters and Violin plots for VenusFP were then generated using ggplot2 v3.3.5 package (69).

#### In vivo T cell proliferation assay

LN cells from pmel-1 TCR transgenic mice were isolated and enriched for CD8+ T cells by magnetic negative selection using CD8+ T cell isolation kit (MACS, Miltenyi). Enriched CD8+ T cells were surface stained and naïve CD8+ T cells were purified by cell sorting (CD8a+, CD62L high, CD44low, CD25 neg). After sorting, naïve CD8+ T cells were labeled with Cell Trace Violet (CTV, Invitrogen). 1×10^6^ naïve CD8+ T cells were adoptively transferred into B16-F10 tumor-bearing mice at day 7 after tumor challenge. *In vivo* proliferation and CD44/CD25 expression of transferred T cells was analyzed by FACS in tumor draining lymph nodes 4 days after adoptive transfer.

#### RNA-seq

Cell suspensions from tumor tissue pooled from 2-4 B16 bearing mice were enriched in immune cells by positive selection with CD45+ biotin magnetic beads (MACS, Miltenyi). Enriched cells were surface stained and 5-20 ×10^3^ intratumoral cDC1s were sorted directly in RLT lysis buffer (Qiagen) containing 2-mercaptoethanol. Immediately after sorting, collected cells were homogenized through vortex and frozen on dry ice before storage at −80°C. Total RNA was extracted with RNAeasy Plus Micro kit (Qiagen). RNA sequencing was performed at VIB Nucleomics Core using SMART-seq v4 pre-amplification followed by single-end sequencing on Illumina NextSeq500. Preprocessing of the RNA-seq data was performed by Trimmomatic v0.39 and quality control by FastQC v0.11.8. Mapping to the reference mouse genome was performed by STAR v2.7.3a and HTSeqCount v0.11.2 was used for counting. Limma v3.42.2 (70) was used to normalize the data. Genes which did not meet the requirement of a count per million (cpm) value larger than 1 in at least 4 samples were filtered. This resulted in an expression table containing 11066 genes. EdgeR v3.28.0 (71) was utilized to perform differential expression analysis. Benjamini-Hochberg correction was used to adjust the p-values for multiple testing. Differentially expressed genes were filtered as genes with a |FC| > 1.5 and adjusted p-value < 0.05. Heatmaps were created using pheatmap v1.0.12 package (72) on log2 normalized and mean centered gene expression data.

#### Gene Set Enrichment Analysis

Over-Representation Analysis (ORA) and Gene Set Enrichment Analysis (GSEA) were performed using *ClusterProfiler* v4.0.5 package (73) in R and Gene Ontology (GO) knowledgebase gene sets. ORA results were considered significant when the q-value was below 0.01. GSEA was performed on pre-ranked mode using as rank metric the signed log10 transformed p-values derived from the differential expression analysis. GSEA was run using the GO:BP database or literature lists of Xbp1-targets and RIDD-targets (47). Results were considered significant when the adjusted p-value was below 0.05.

### QUANTIFICATION AND STATISTICAL ANALYSIS

No statistical methods were used to predetermine sample size. The experiments were not randomized, and the investigators were not blinded to allocation during experiments and outcome assessment. Statistical analysis was conducted using GraphPad Prism software (v9.1.2). Results are presented as mean ± SEM. Two groups were compared using two tailed t-test for normal distributed data (Shapiro-Wilk test) or using a non-parametric two-tailed Mann-Whitney test as indicated in figure legends. Multiple groups were compared using one-way ANOVA with Tukey post-test. A p-value < 0.05 was considered statistically significant.

### STUDY APPROVAL

All animal procedures were performed in accordance with institutional guidelines for animal care of the Fundación Ciencia y Vida, the Faculty of Medicine, University of Chile and the VIB, Belgium, and were approved by the local ethics committee.

## ABBREVIATIONS

BM: bone marrow
cDC: conventional DC
cDC1: type 1conventional DC (XCR1^+^ DC)
cDC2: type 2conventional DC (CD11b^+^ DC)
DC: dendritic cell
DEG: Differentially expressed gene
ER: endoplasmic reticulum
ERAI: ER stress-activated indicator
Flt3L: FMS-related tyrosine kinase 3 ligand
FP: fluorescent protein
GSEA: Gene set enrichment analysis
IRE1: inositol-requiring enzyme 1 alpha
KO: Knock-out
MdC: Myeloid derived Cell
RIDD: regulated IRE1-dependent decay
ROS: reactive oxygen species
TAM: tumor-associated macrophages
TCR: T cell receptor
TdLN: Tumor draining lymph node
UPR: unfolded protein response
XBP1s: spliced XBP1
XBP1u: unspliced XBP1

## Author contributions

F.F, MR.B, S.J and F.O designed the research; F.F, S.R, S.G, C.F, D.F did the experiments. F.F, S.R, S.J and F.O analyzed the results; C.DN and C.M helped with RNA-seq data analysis; D.Fe provided technical assistance and experimental expertise, T.I and A.L provided critical reagents. F.F. and F.O wrote the manuscript

## Acknowledgements

We thank Dr Laurie H. Glimcher (Dana-Farber Cancer Institute) for *Xbp1*^fl/fl^ mice; Dr Matthew Krummel (UCSF) for the B78ChOVA cell line; Dr. Juan Carlos Zuñiga-Pflucker (Sunnybrook Research Institute) for the OP9-DL1 cell line; the VIB nucleomics facility for doing the RNA seq experiments and facilities at Universidad de Chile and Fundación Ciencia & Vida. We thank members of the immunology and immunology and cellular stress laboratories for critical support.

This work was funded by an International Research Scholar grant from HHMI (HHMI#55008744, FO); FONDECYT grant No 1200793 (FO); FONDECYT grant No 1191438 (MR.B); CONICYT/FONDEQUIP/EQM140016; FONDECYT grant No 1212070 (AL); ANID grant FB210008 (MR.B and AL); CONICYT-PFCHA/DoctoradoNacional/2017-21170366 (F.F). The work in Belgium was funded by Stichting tegen Kanker (2014/283), FWO-EOS (ID 30837538) and ERC-CoG (ID 819314) (S.J.).

## SUPPLEMENTARY FIGURE LEGENDS

**Supplementary Figure 1. Immune analysis of melanoma tumors derived from ERAI mice.**

**(A)** t-SNE map as in figure 1a. Color gradient shows the expression of the indicated marker. **(B)** Quantification of VenusFP signal from manually gated immune populations from MC-38 bearing WT and ERAI mice. **** p < 0.0001, ANOVA and Tukey post-test between ERAI mice. n=3 (WT) or 5 (ERAI) mice per group, representative of two independent experiments.(C) Quantification of VenusFP signal of tumor- and spleen-cDC1s from MC-38 bearing WT and ERAI mice. **** p<0.0001, t-test between ERAI mice. n=3 (WT) or 5 (ERAI), representative of two independent experiments.

**Supplementary Figure 2. Gating strategy for tumor associated cDCs.**

**(A)** Gating strategy for identification of immune infiltrated populations in tumors. Representative plots from B16 melanoma tumor. **(B)** Gating strategy for identification of migratory (mig) and resident (res) cDC1s and cDC2s in tumor draining lymph node. Representative plots from B16 melanoma TdLN.

**Supplementary Figure 3. Tumor growth and T cell infiltration in MC-38 bearing XBP1^ΔDC^ mice.**

XBP1^WT^ and XBP1^ΔDC^ mice were implanted with MC-38 tumors. **(A-C)** Tumor volume curves (A), tumor volumes (B) at end point and tumor weight (C) at end point. n=12-13 mice per group. Pooled data from two independent experiments.

**Supplementary Figure 4. Tumor immune cell analysis of XBP1^ΔDC^ mice.**

**(A)** Related to figure 3A. CD40 and PD-L1 expression by TdLN cDCs. n=7 mice per group, pooled data from two independent experiments, mean ± s.e.m. **(B)** Related to figure 3B. Dot plots of IL-12 intracellular expression by *in vitro* generated cDC1s stimulated with B78-ChOVA lysates. **(C)** Related to figure 3C-D. Fluorescence minus one (FMO) and tumor-free WT mouse as negative controls for tetramer staining. **(D-E)** XBP1^WT^ and XBP1^ΔDC^ mice were implanted with B16-F10 tumors. Seven days after implantation, CD8^+^ naïve T cells isolated from pmel1 transgenic mice were labeled with Cell Trace Violet and were adoptively transferred into tumor-bearing mice. Four days later, transferred cell proliferation and CD44/CD25 expression was quantified by FACS. n=8 mice (XBP1^WT^) or 15 mice (XBP1^ΔDC^), data pooled from two independent experiments, mean ± s.e.m. **(F-G)** CD8+ T cell (F) and CD4+ T cell (G) frequencies and profiles in MC-38 bearing XBP1^WT^ and XBP1^ΔDC^ mice. n=11-13, pooled data from two independent experiments, mean ± s.e.m. **(H)** Related to figure 3H-I. Representative histograms of different markers associated with terminal exhausted CD8^+^ T cells. Gated on CD3^+^CD8^+^PD1^+^TCF1^+^ or CD3^+^CD8^+^PD1^+^TIM3^+^ as shown in figure 3H.

**Supplementary Figure 5. IRE1/XBP1s double deficient cDC1s are unable to activate RIDD.**

**(A)** PCR analysis of XBP1 splicing in intratumoral cDC1s isolated from B16-bearing XBP1^WT^/IRE1^WT^ vs XBP1^ΔDC^/IRE1^trunc^DC mice. Each lane represents different mice. XBP1u: XBP1 unspliced; Xbp1s: Xbp1 spliced; Actb: beta actin. **(B)** CD11c expression by intratumoral cDC1, cDC2 and TAMs from B16-bearing XBP1^WT^/IRE1^WT^ vs XBP1^ΔDC^/IRE1^trunc^DC mice. n=4 mice per group, mean ± s.e.m. **(C)** CD11c expression by cDC subsets in the TdLN from B16-bearing XBP1^WT^/IRE1^WT^ vs XBP1^ΔDC^/IRE1^trunc^DC mice. *p<0.05, two-tailed Mann-Whitney test. n=4 mice per group, mean ± s.e.m.

**Supplementary Figure 6. RNAseq analysis of WT, XBP1 deficient WT vs IRE1/XBP1 double deficient cDC1s.**

**(A)** Heatmap of genes with an adjusted p-value < 0.05 but with a fold change (FC) below the 1.5 threshold. Table summarize log2FC and adj p-values for WT vs XBP1 and WT vs IRE1/XBP1 deficient cDC1s. **(B)** Related to figure 5F-G. GSEA using XBP1s- and RIDD-target gene sets from literature (So et al, 2012, Cell Metabolism). **(C)** GSEA of the “triglyceride biosynthetic process” gene set (GO: 0019432) in WT vs XBP1^ΔDC^ cDC1s (left) or WT vs XBP1^ΔDC^/IRE1^trunc^DC cDC1s (right) showing not statistically significant enrichment (q-value > 0.05). **(D)** Normalized expression (z-scores) for genes of the “triglyceride biosynthetic process” gene set (GO: 0019432) in WT, XBP1^ΔDC^, or XBP1^ΔDC^/IRE1^tr^u^nc^DC cDC1s.

