## Supplemental files for "Nuanced role for dendritic cell intrinsic IRE1 RNase in the regulation of antitumor adaptive immunity"

This file contains:

- One supplementary reagents and tools table
- Supplementary Figures 1-6.

**Reagents and Tools table**

| REAGENT or RESOURCE | SOURCE | IDENTIFIER |
|---|---|---|
| Antibodies | | |
| B220 BV510 | Biolegend | clone RA3-6B2; #cat 103222; RRID:AB_313005 |
| B220 PE-Cy5 | Biolegend | clone RA3-6B2; #cat 103209; RRID:AB_312994 |
| CD11b BV785 | Biolegend | clone M1/70; #cat 101243; RRID:AB_2561373 |
| CD11c APC | Thermo Fisher Scientific | clone N418; #cat 17-0114-82; RRID:AB_469346 |
| CD11c BV711 | Biolegend | clone N418; #cat 117349; RRID:AB_2563905 |
| CD16/32 (FcBlock) - | Biolegend | clone 93; #cat 101302; RRID:AB_312801 |
| CD18 PE | BD Biosciences | clone C71/16; #cat 553293; RRID:AB_394762 |
| CD19 BUV737 | BD Biosciences | clone 1D3; #cat 564296; RRID:AB_2716855 |
| CD25 PE | Tonbo Bioscience | clone PC61.5; #cat 50-0251; RRID:AB_2621757 |
| CD26 PE-Cy7 | Biolegend | clone H194-112; #cat 137809; RRID:AB_2564311 |
| CD274 (PD-L1) PE | Thermo Fisher Scientific | clone MIH5; #cat 12-5982-82; RRID:AB_466089 |
| CD3 BV711 | Biolegend | clone 17A2; #cat 100241; RRID:AB_2563945 |
| CD39 SuperBright600 | Thermo Fisher Scientific | clone 24DMS1; #cat 63-0391-82; RRID:AB_2717037 |
| CD3ε BV510 | Biolegend | clone 145-2C11; #cat 100353; RRID:AB_2565879 |
| CD3ε PE-Cy5 | Biolegend | clone 145-2C11; #cat 100309; RRID:AB_312674 |
| CD4 APC-Cy7 | Tonbo Bioscience | clone GK1.5; #cat 25-0041; RRID:AB_2904484 |
| CD4 PE/Dazzle 594 | Biolegend | clone GK1.5; #cat 100455; RRID:AB_2565844 |

| | | |
|---|---|---|
| CD40 APC | Biolegend | clone 3/23.; #cat 124611; RRID:AB_1134081 |
| CD44 PerCP | Biolegend | clone IM7; #cat 103036; RRID:AB_10645506 |
| CD44 FITC | Biolegend | clone IM7; #cat 103006; RRID:AB_312957 |
| CD45 BV785 | Biolegend | clone 30-F11 ; #cat 103149; RRID:AB_2564590 |
| CD45 Alexa Fluor 700 | Biolegend | clone 30-F11; #cat 103127; RRID:AB_493714 |
| CD45 BUV395 | BD Biosciences | clone 30-F11; #cat 564279; RRID:AB_2651134 |
| CD45 BV650 | Biolegend | clone 30-F11; #cat 103151; RRID:AB_2565884 |
| CD45.1 APC-Cy7 | Biolegend | clone A20; #cat 110715; RRID:AB_313504 |
| CD62L PE-Cy7 | Biolegend | clone MEL-14 ; #cat 104418; RRID:AB_313103 |
| CD64 BV711 | Biolegend | clone X54-5/7.1 ; #cat 139311; RRID:AB_2563846 |
| CD64 PE/Dazzle 594 | Biolegend | clone X54-5/7.1; #cat 139319; RRID:AB_2566558 |
| CD64 PerCP-Cy5.5 | Biolegend | clone X54-5/7.1; #cat 139307; RRID:AB_2561962 |
| CD8a APC-Cy7 | Tonbo Bioscience | clone 53-6.7; #cat 25-0081; RRID:AB_2621623 |
| CD8a BV650 | Biolegend | clone 53-6.7; #cat 100742; RRID:AB_2563056 |
| CD8a eFluor 450 | Thermo Fisher Scientific | clone 53-6.7; #cat 48-0081-80; RRID:AB_1272235 |
| F4/80 Biotin | Biolegend | clone BM8; #cat 123105; RRID:AB_893499 |
| FOXP3 PE-Cy7 | Thermo Fisher Scientific | clone FJK-16s; #cat 25-5773-82; RRID:AB_891552 |
| Granzyme B PE Texas RED | Thermo Fisher Scientific | clone GB11; #cat GRB17; RRID:AB_2536540 |

| | | |
|---|---|---|
| I-A/I-E (MHC-II) APC-Cy7 | Biolegend | clone M5/114.15.2; #cat 107627; RRID:AB_1659252 |
| IFN-γ PE | Thermo Fisher Scientific | clone XMG1.2; #cat 12-7311-82; RRID:AB_466193 |
| IL-12p40 eFluor660 | Thermo Fisher Scientific | clone C17.8; #cat 50-7123-80; RRID:AB_11218284 |
| IL-2 PE-Cy7 | Thermo Fisher Scientific | clone JES6-5H4; #cat 25-7021-82; RRID:AB_1235004 |
| Ly6C BV605 | Biolegend | clone HK1.4; #cat 128035; RRID:AB_2562352 |
| Ly6G Alexa Fluor 700 | Biolegend | clone 1A8; #cat 127622; RRID:AB_10643269 |
| NK1.1 PerCP-Cy5.5 | Biolegend | clone PK136; #cat 108727; RRID:AB_2132706 |
| PD1 PE | Biolegend | clone 29F.1A12; #cat 135206; RRID:AB_1877231 |
| PD1 BV421 | Biolegend | clone 29F.1A12; #cat 135217; RRID:AB_10900085 |
| TCF1 Alexa Fluor 488 | Cell Signaling | clone C63D9; #cat CS.6444S; RRID:AB_2797627 |
| TIM3 PECy7 | Biolegend | clone RMT3-23; #cat 119716; RRID:AB_2571933 |
| TNF-α APC | Biolegend | clone MP6-XT22; #cat 506307; RRID:AB_315428 |
| TOX APC | Miltenyi Biotec | clone REA473; #cat 130-118-335; RRID:AB_2751485 |
| XCR1 BV650 | Biolegend | clone ZET; #cat 148220; RRID:AB_2566410 |
| XCR1 PE | Biolegend | clone ZET; #cat 148204; RRID:AB_2563843 |
| **Bacterial and virus strains** | | |
| **Biological samples** | | |
| Mouse tumor draining lymph nodes (inguinal) | This paper | N/A |
| Mouse tumor | This paper | N/A |
| Mouse spleen | This paper | N/A |
| **Chemicals, peptides, and recombinant proteins** | | |
| Brefeldin A | Cayman Chemical | #cat 11861 |
| PMA | Sigma | #cat P8139 |

| | | |
|---|---|---|
| Ionomycin | Sigma | #cat I0634 |
| Critical commercial assays | | |
| CD45 MicroBeads, mouse | Miltenyi | RRID:AB_2877061 |
| CD8a+ T Cell Isolation Kit, mouse | Miltenyi | #cat 130-104-075 |
| LS columns | Miltenyi | #cat 130-042-401 |
| BD Cytofix/Cytoperm with GolgiPlug | BD Biosciences | RRID:AB_2869013 |
| Foxp3 / Transcription Factor Staining Buffer Set | eBioscience | #cat 00-5523-00 |
| RNeasy Plus Micro Kit | Qiagen | #cat 74034 |
| M-MLV Reverse Transcriptase | Invitrogen | #cat 28025-013 |
| iTAg Tetramer/PE - H-2 Kb OVA (SIINFEKL) | MBL | #cat T03000 |
| CellTrace™ Violet Cell Proliferation Kit | Thermo Fisher Scientific | #cat C34557 |
| Deposited data | | |
| RNA-seq data | This paper | GEO: GSE195439 |
| Experimental models: Cell lines | | |
| B16-F10 | ATCC | RRID:CVCL_0159 |
| B16-FLT3L | (Guo-Ping *et al.*, 1999) | N/A |
| B78-ChOVA | (Broz *et al.*, 2014) | N/A |
| OP9-DL1 | (Schmitt and Zúñiga-Pflücker, 2002) | RRID:CVCL_B218 |
| Experimental models: Organisms/strains | | |
| ERAI: C57BL/6J-Tg(CAG-XBP1*/venus)#Miur/MiurRbrc | (Iwawaki *et al.*, 2004) | RRID:IMSR_RBRC01099 |
| CD11c-Cre: B6.Cg-Tg(Itgax-cre)1-1Reiz/J | The Jackson Laboratory | RRID:IMSR_JAX:008068 |
| IRE1 fl/fl: B6;129S4-Ern1<tm2.1Tiw> | (Iwawaki *et al.*, 2009) | RRID:IMSR_RBRC05515 |
| XBP1 fl/fl: Xbp1tm2Glm | (Lee *et al.*, 2008) | RRID:MGI:6273536 |
| Pmel-1: B6.Cg-Thy1a/Cy Tg(TcraTcrb)8Rest/J | The Jackson Laboratory | RRID:IMSR_JAX:005023 |
| Oligonucleotides | | |
| Primer: Splicing assay Xbp1 Forward: ACACGCTTGGGAATGGACAC | (Osorio *et al.*, 2014) | N/A |
| Primer: Splicing assay Xbp1 Reverse: CCATGGGAAGATGTTCTGGG | (Osorio *et al.*, 2014) | N/A |
| Primer: Actb Forward (Fig 1G): CTAAGGCCAACCGTGAAAAG | This Paper | N/A |
| Primer: Actb Reverse (Fig 1G): TTGCTGATCCACATCTGCTG | This Paper | N/A |
| Primer: Actb Forward (Fig 4B and S5A): GTGACGTTGACATCCGTAAAGA | This Paper | N/A |
| Primer: Actb Reverse (Fig 4B and S5A): GCCGGACTCATCGTACTCC | This Paper | N/A |
| Recombinant DNA | | |
| Software and algorithms | | |
| FlowJo™ Software v10 | BD Biosciences | https://www.flowjo.com/solutions/flowjo; RRID:SCR_008520 |

| | | |
|---|---|---|
| GraphPad Prism v9 | GraphPad | https://www.graphpad.com/; RRID:SCR_002798 |
| Trimmomatic v0.39 | Usadellab | RRID:SCR_011848 |
| FastQC v0.11.8 | Babraham Bioinformatics | RRID:SCR_014583 |
| STAR v2.7.3a | STAR | RRID:SCR_004463 |
| HTSeqCount v0.11.2 | HTSeq | RRID:SCR_005514 |
| R Studio v2021.09.0 Build 351 | R Studio | https://rstudio.com/; RRID:SCR_000432 |
| R v4.1.1 | R Core Team (2020) | http://www.r-project.org/; RRID:SCR_001905 |
| R package: CytoML v2.4.0 | (Finak, Jiang and Gottardo, 2018) | https://bioconductor.org/packages/release/bioc/html/CytoML.html |
| R package: FlowWorkspace v4.4.0 | (Finak and Jiang, 2021) | RRID:SCR_001155 |
| R package: FlowCore v2.4.0 | (Ellis *et al.*, 2021) | RRID:SCR_002205 |
| R package: Rtsne v0.15 | (Krijthe, 2015) | RRID:SCR_016342 |
| R package: DBSCAN v1.1.8 | (Hahsler, Piekenbrock and Doran, 2019) | https://cran.r-project.org/web/packages/dbscan/index.html |
| R package: Limma v3.42.2 | (Ritchie *et al.*, 2015) | RRID:SCR_010943 |
| R package: EdgeR v3.28.0 | (Robinson, McCarthy and Smyth, 2010) | RRID:SCR_012802 |
| R package: pheatmap v1.0.12 | (Kolde, 2019) | RRID:SCR_016418 |
| R package: ggplot2 v3.3.5 | (Wickham, 2016) | RRID:SCR_014601 |
| R package: ClusterProfiler v4.0.5 | (Wu *et al.*, 2021) | RRID:SCR_016884 |
| Other | | |

### Supplementary Figure 1

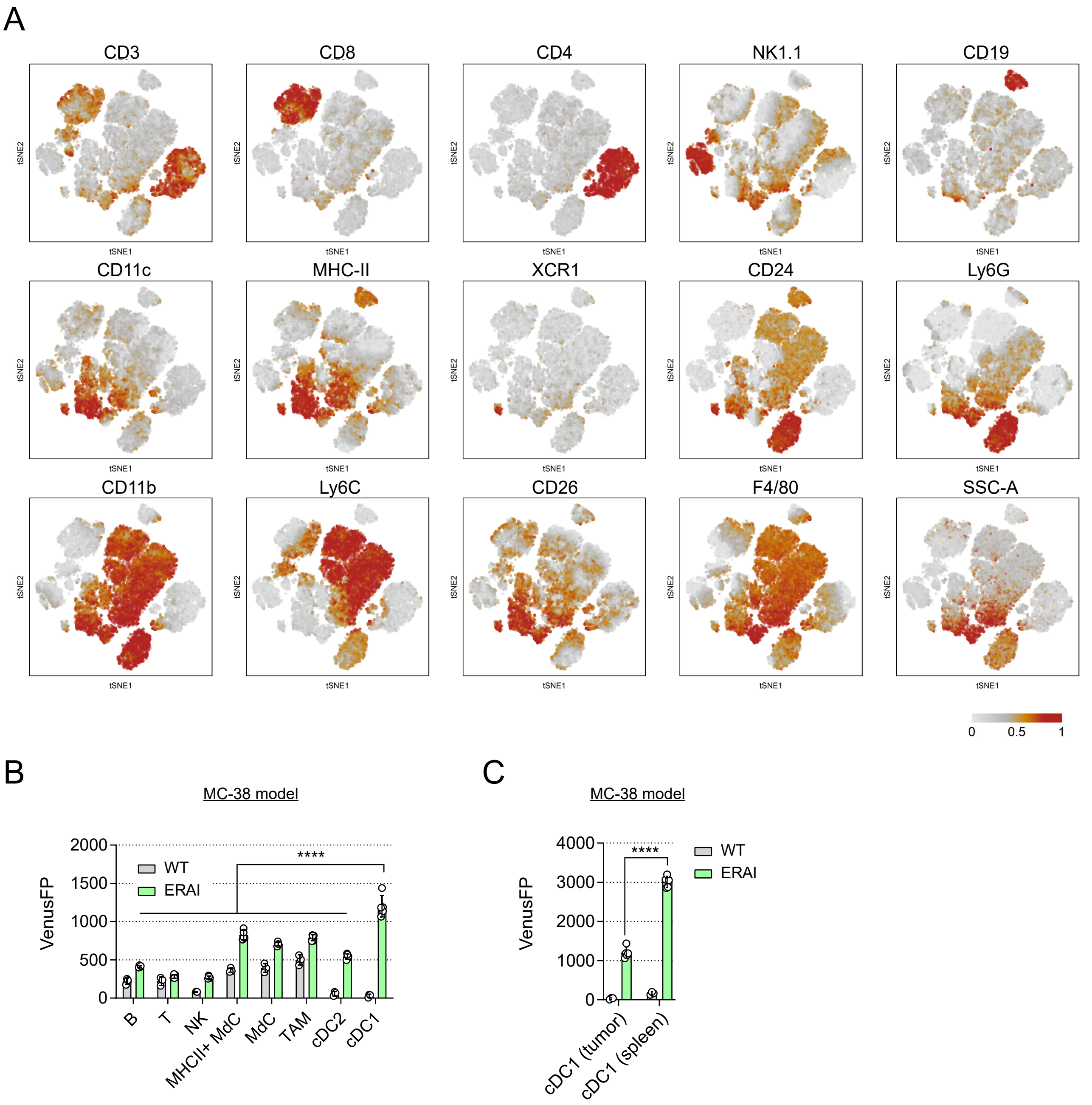

### Supplementary Figure 2

## A

Tumor gating

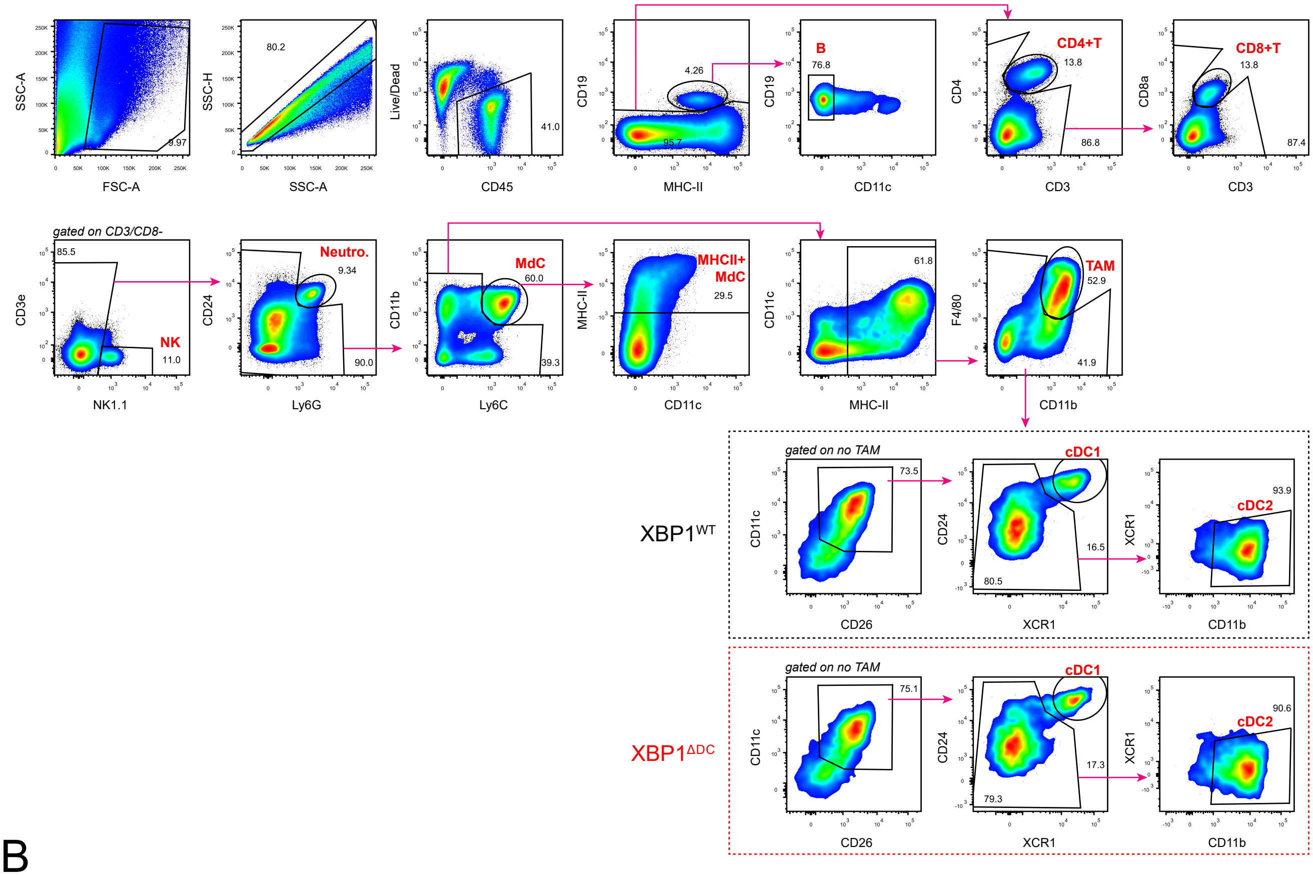

## B

Tumor draining lymph node gating

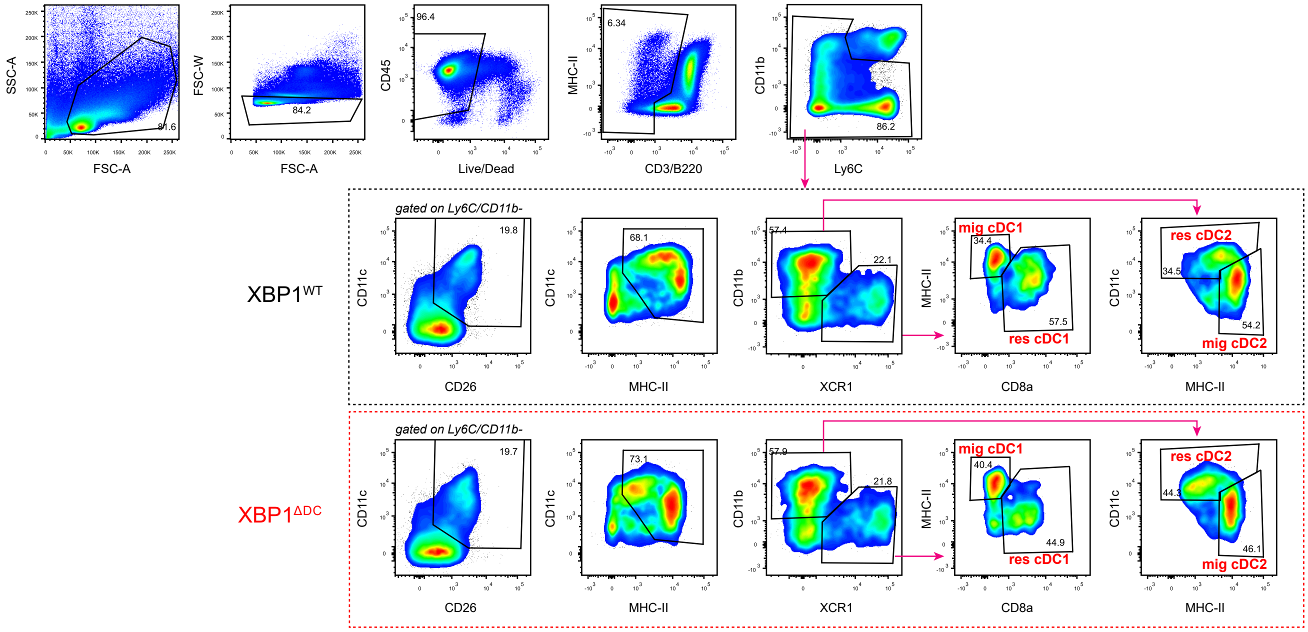

### Supplementary Figure 3

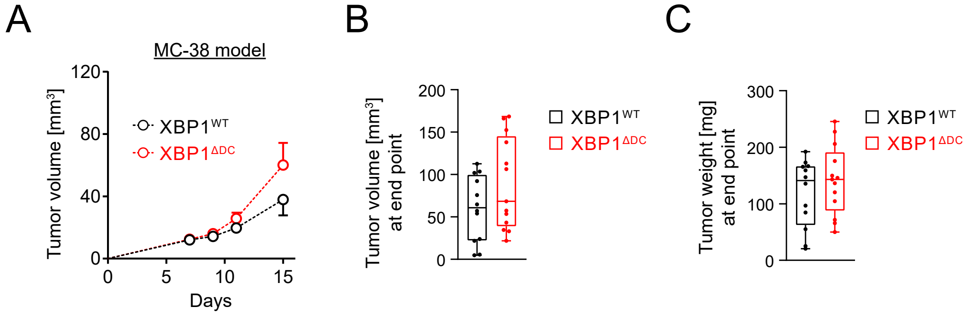

### Supplementary Figure 4

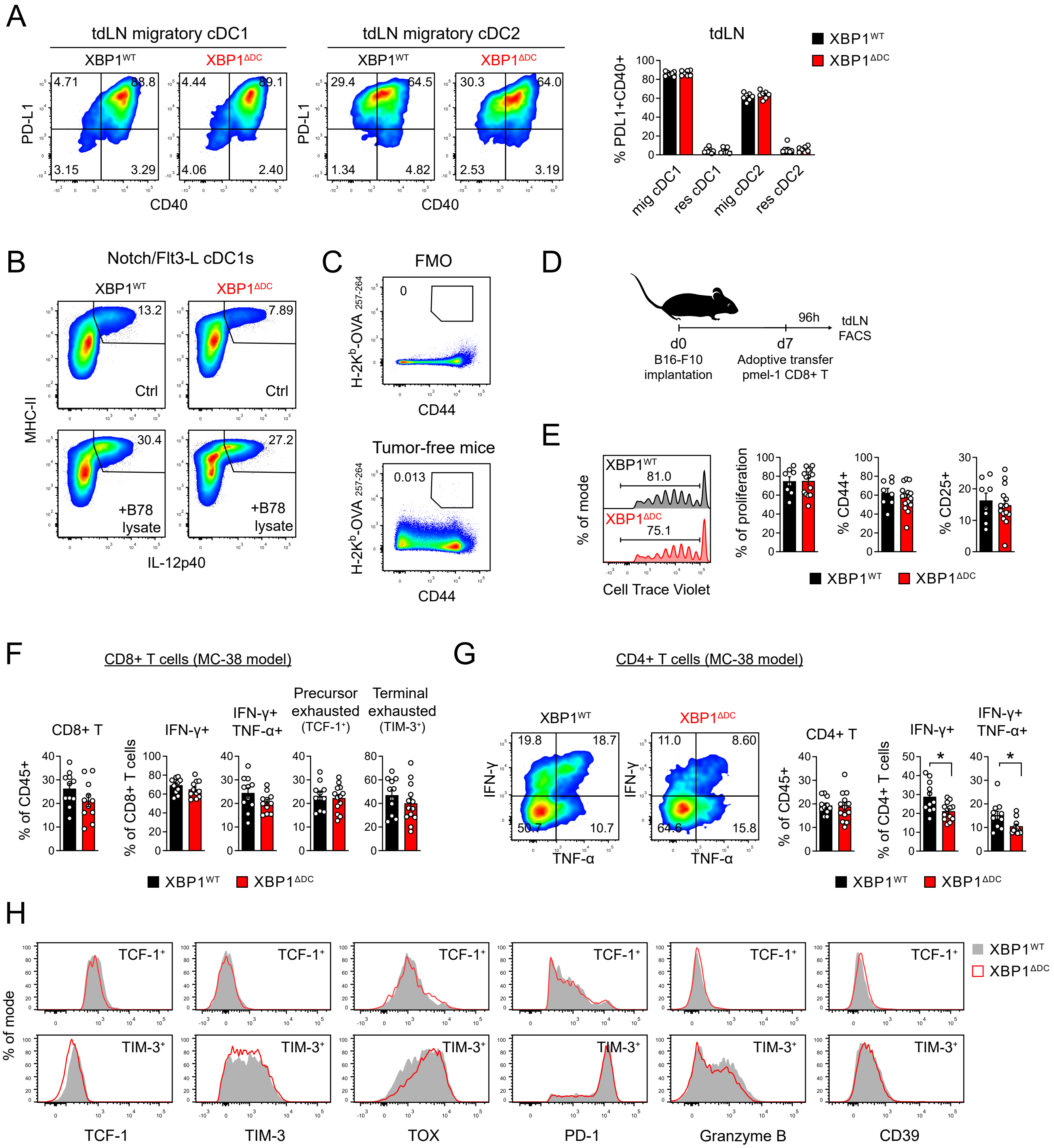

### Supplementary Figure 5

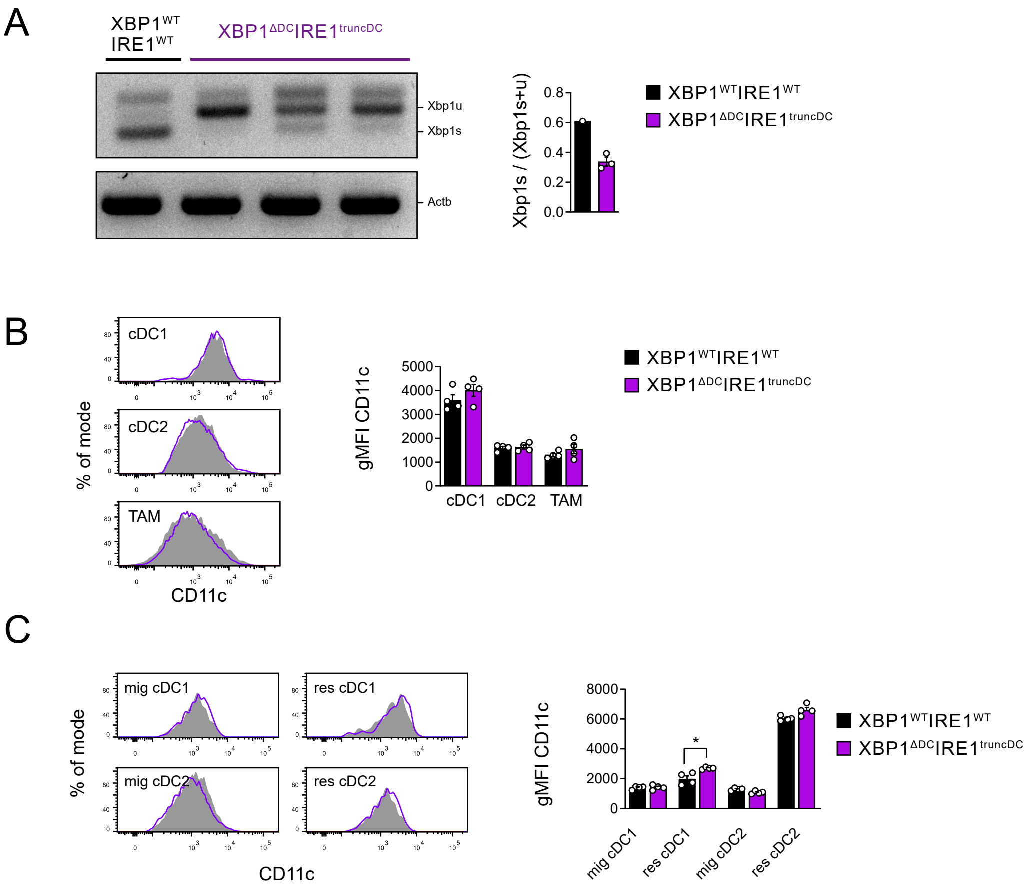

### Supplementary Figure 6

A

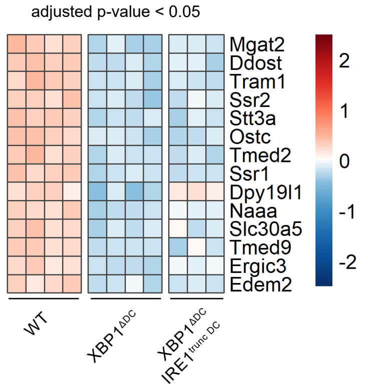

| | WT vs XBP1ΔDC | | WT vs XBP1ΔDC IRE1truncDC | |
|---|---|---|---|---|
| gene | log2FC | adj p-value | log2FC | adj p-value |
| Ddost | -0.571 | 0.015 | -0.529 | 0.056 |
| Dpy19l1 | -0.520 | 0.039 | -0.080 | 0.839 |
| Edem2 | -0.424 | 0.048 | -0.409 | 0.128 |
| Ergic3 | -0.460 | 0.016 | -0.284 | 0.234 |
| Mgat2 | -0.577 | 0.016 | -0.470 | 0.128 |
| Naaa | -0.512 | 0.013 | -0.328 | 0.167 |
| Ostc | -0.532 | 0.016 | -0.556 | 0.041 |
| Slc30a5 | -0.494 | 0.036 | -0.372 | 0.214 |
| Ssr1 | -0.523 | 0.010 | -0.517 | 0.027 |
| Ssr2 | -0.537 | 0.016 | -0.457 | 0.108 |
| Stt3a | -0.535 | 0.016 | -0.527 | 0.045 |
| Tmed2 | -0.525 | 0.016 | -0.548 | 0.040 |
| Tmed9 | -0.485 | 0.039 | -0.455 | 0.128 |
| Tram1 | -0.540 | 0.016 | -0.505 | 0.058 |

B

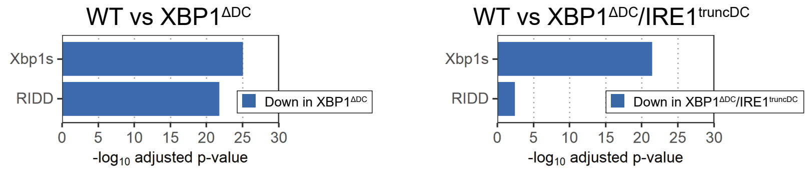

C

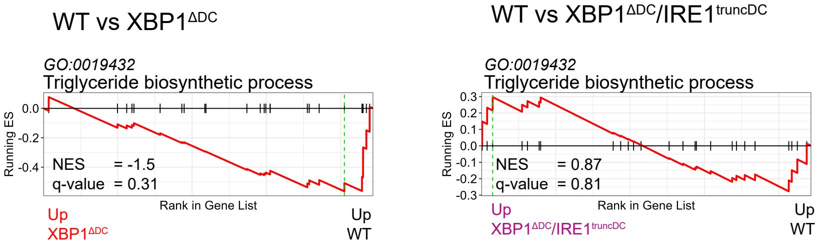

D

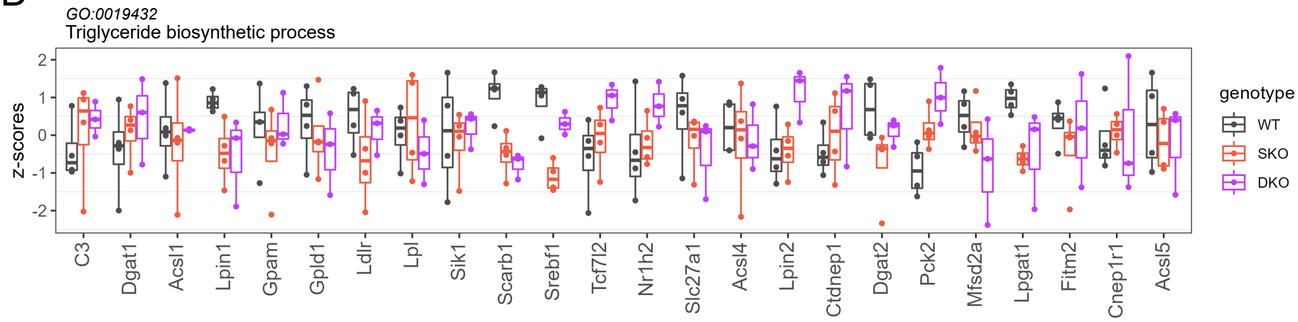
